# The Initiator Caspase Dronc Drives Compensatory Proliferation of Apoptosis-Resistant Cells During Epithelial Tissue Regeneration After Ionizing Radiation

**DOI:** 10.1101/2024.09.01.610661

**Authors:** Tslil Braun, Naama Afgin, Lena Sapozhnikov, Ehud Sivan, Andreas Bergmann, Luis Alberto Baena-Lopez, Keren Yacobi-Sharon, Eli Arama

## Abstract

Caspases, well-known for their role in executing apoptosis, also participate in various non-apoptotic processes. Despite this, their involvement in promoting compensatory proliferation - a key aspect of tissue regeneration following extensive cell death - has been a subject of ongoing ambiguity. In our study, we investigate compensatory proliferation in the *Drosophila* wing imaginal disc following ionizing radiation, a model epithelial tissue that has been a pioneering system for studying this regenerative response. Using a delayed genetic reporter to monitor the activity of the initiator caspase-2/9 ortholog, Dronc, we identified two populations of apoptosis-resistant epithelial cells involved in compensatory proliferation: those that activate Dronc (termed DARE cells) and those that do not (NARE cells). We show that DARE cells pass their apoptosis-resistance trait to their daughter cells, suggesting a molecular memory. We demonstrate that Dronc in DARE cells, but not the apoptosome adapter Dark and the effector caspases, promotes compensatory proliferation both within these cells and in NARE cells through a non-cell-autonomous mechanism. We found that Myo1D, an unconventional myosin interacting with Dronc, is essential for the survival of DARE cells by preventing the lethal activation of effector caspases and subsequent apoptosis. In contrast, Myo7A/Crinkled, another unconventional myosin that interacts with Dronc, promotes effector caspase activation in DARE cells. We demonstrate that the TNFR>JNK signaling pathway in DARE cells directly regulates their proliferation, which in turn influences NARE cell proliferation. Consequently, we show that maintaining proliferative homeostasis between DARE and NARE cells is vital for balanced tissue regeneration. Given the widespread use of ionizing irradiation in cancer treatment and prevention, our findings have potential implications for understanding treatment-resistant cells and cancer recurrence.

## Introduction

Programmed cell death (PCD) is a genetically regulated process that eliminates unwanted or potentially dangerous cells during development and homeostasis in multicellular organisms. ^1–3^ Disruption of PCD is linked to various diseases, including cancer and neurodegenerative disorders. ^1,4–6^ Apoptosis, the most prevalent form of PCD during animal development, is characterized by a conserved sequence of morphological and cellular events. ^7,8^ Biochemically, apoptosis is marked by the activation of caspases, a unique family of cysteine proteases. ^9,10^ Caspases are crucial for both the signaling and execution of apoptosis, and their activation is tightly regulated by various activating and inhibitory proteins. ^10–14^ Synthesized as inactive pro-enzymes, caspases undergo a precisely controlled proteolytic cascade to become active and to activate each other. ^10,11,15–17^ Initiator caspases, such as caspase-9 (intrinsic pathway) and caspase-8 (extrinsic pathway), cleave and activate effector caspases like caspase-3 and caspase-7, ^18,19^ which orchestrate cell death by cleaving numerous cellular proteins. ^20,21^ Inadvertent activation of caspases is prevented by inhibitory proteins such as the Inhibitor of Apoptosis proteins (IAPs), which bind to and inhibit caspases. ^22–25^

The *Drosophila* genome encodes seven caspases, with the initiator caspase-9 ortholog Dronc and the effector caspases 3 and 7 orthologs, Drice and Dcp-1, mediating most apoptotic events during development and stress. ^26–30^ Dronc, similar to caspase-9, is activated on the apoptosome through binding to the *Drosophila* Apaf-1 related killer (Dark) adapter protein and subsequently activates Drice and Dcp-1. ^29,31–37^ In living cells, the major *Drosophila* IAP, Diap1, inhibits Dronc and the effector caspases. ^22,38–40^ During apoptosis, pro-apoptotic Reaper family proteins bind to Diap1, relieving its inhibition of caspases. ^22,25,41–43^

Despite their well-established role in apoptosis, numerous non-lethal <u>c</u>aspase-<u>d</u>ependent cellular <u>p</u>rocesses (CDPs) have been identified across various tissues and organisms. While the molecular mechanisms governing apoptosis and caspase activation are well-understood, research into CDPs has progressed more slowly due to the complexities of studying these processes in specific cell types and tissues *in vivo*. Nevertheless, over the past two decades, CDPs have garnered significant attention due to their important biological and pathological implications. ^44–53^

Compensatory proliferation is a regenerative response to extensive cell death due to trauma. Although the involvement of caspases has been proposed, the molecular details remained unclear. ^54–57^ Early evidence of this phenomenon was observed in *Drosophila* wing imaginal discs (WIDs), which regenerate after high-dose ionizing radiation (IR), presumably by increasing the proliferation of surviving cells. ^58–60^ Despite this, identifying and tracking the cells responsible for this regeneration has been challenging, inevitably leading to confusion with other tissue regeneration mechanisms. One such mechanism, known as apoptosis-induced proliferation (AiP), has particularly complicated the understanding of compensatory proliferation. ^54,57,61–70^ AiP occurs when cells, induced to undergo apoptosis by pro-apoptotic gene overexpression or IR, are kept alive by the ectopic expression of the baculovirus P35 protein, a pan-effector caspase inhibitor. These ‘undead’ cells activate Dronc but cannot undergo apoptosis, leading to hyperproliferation and overgrowth in neighboring wild-type (WT) cells. ^61,63–65^ This led to the hypothesis that AiP might mediate compensatory proliferation via non-cell autonomous signals from dying cells. ^42,60,71,72^ However, AiP is a non-physiological phenomenon of ‘undead’ cells rather than dying cells. Moreover, compensatory proliferation continues even when key AiP-associated signals, such as Wingless/Wnt and Dpp/BMP, are disrupted.^73^ This distinction suggests that AiP and compensatory proliferation are separate phenomena, ^54^ though they may share some mechanistic features.

Here, we show that compensatory growth of the WID after IR is mediated by the balanced proliferation of two cell populations: one that activates Dronc, referred to as DARE cells (<u>D</u>ronc-activating <u>a</u>poptosis resistant <u>e</u>pithelial cells), and another that does not activate caspases, termed NARE cells (<u>n</u>on-Dronc-activating <u>a</u>poptosis resistant <u>e</u>pithelial). Tracking DARE cell lineage reveal that multiple cells appear at 24 hours post-irradiation (hpi) and undergo extensive proliferation over the subsequent 24 hours, leading to tissue regeneration. Re-irradiation of DARE cell clones at 48 hpi results in significantly fewer dying cells 4 hours later compared to the initial irradiation, suggesting a molecular memory of the resistance trait. Ablation of DARE cells, as well as downregulation of Dronc in these cells, significantly impairs both DARE and NARE cell proliferation and tissue regeneration, resulting in smaller WIDs at 48 hpi. In contrast, inactivation of the apoptosome adapter Dark or the effector caspases in DARE cells has no detectable effect on the regeneration of this tissue. Knockdown of *myo1D* in DARE cells - a gene encoding an unconventional myosin that sequesters Dronc to the basal side of the cell membrane during AiP - results in significant activation of effector caspases and subsequent DARE cell death. This reveals a mechanism that regulates effector caspase activation to ensure the survival of these cells despite the presence of active Dronc. In contrast, Myo7A/Crinkled, another unconventional myosin that interacts with Dronc, promotes effector caspase activation in DARE cells, which likely at least partially explains why not all DARE cells survive. Interestingly, we provide evidence for the involvement of the TNF receptors and JNK in the regulation of DARE cell proliferation, which, in turn, coordinates the proliferation of NARE cells. Finally, we demonstrate that the proliferative homeostasis between DARE and NARE cells, crucial for balanced tissue regeneration, is maintained through cell competition-like mechanisms. Specifically, a higher proliferative rate or fitness of one cell population results in the outcompeting of the other population.

## Results

### Delayed Reporting of the Initiator Caspase Dronc Activity Highlights Apoptosis-Resistant Epithelial Cells

To explore the mechanisms behind compensatory proliferation in the *Drosophila* wing imaginal disc (WID), we first established a timeline of tissue regeneration by monitoring apoptotic cell counts and changes in tissue area at various time points following exposure to 20 Gy of X-radiation. As detailed below, we focused on three specific time points: 4, 24, and 48 hours post-irradiation (hpi). These time points correspond to the approximate onset of apoptosis (4 hpi), a significant accumulation of dying cells and a decrease in WID size (24 hpi), and the post-regeneration phase when almost all dying cells are cleared and normal tissue size is restored (48 hpi). Apoptotic cells were identified through immunostaining with the anti-cleaved Dcp-1 antibody, which specifically detects the cleaved/activated forms of the effector caspases Drice and Dcp-1 ^29^ (jointly referred to here as cDcp-1). As previously reported, ^29,74^ numerous apoptotic cells are detected as early as 4 hpi, with a significant increase observed around 24 hpi (Figures 1A and 1B). Consequently, measurements of the WID area showed a significant reduction in size at both 4 and 24 hpi (Figures 1A and 1C). At 48 hpi, the number of apoptotic cells decreases dramatically, and the WID area returns to normal size (Figures 1A-1C). These findings suggest that most cell death in the irradiated WID occurs within the first ∼24 hpi, while extensive tissue regeneration, likely driven by the compensatory proliferation of apoptosis-resistant cells, predominantly takes place during the subsequent ∼24 hours.

**Figure 1.**
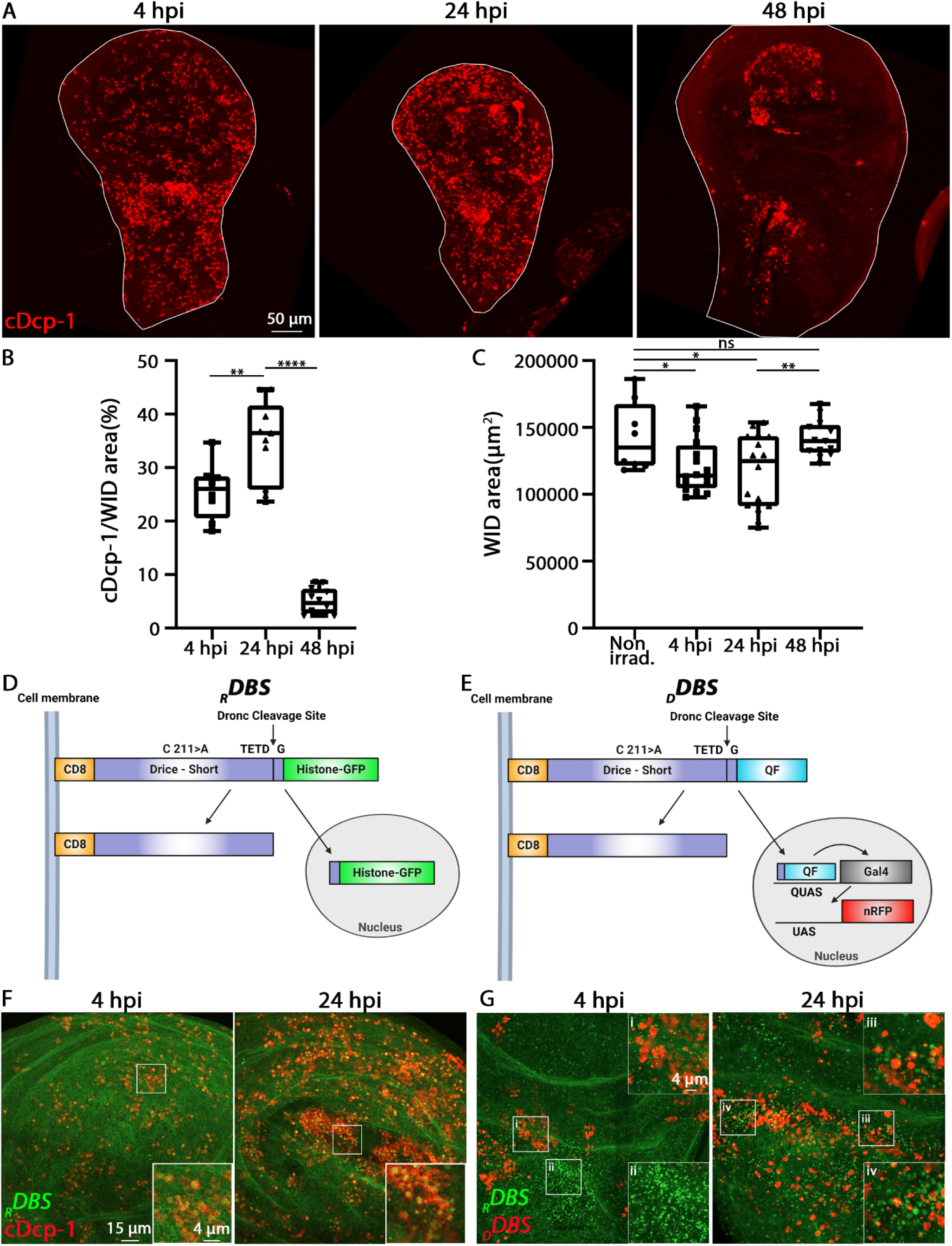
Detection of irradiation-induced apoptosis-resistant cells using a delayed reporter of Dronc activity. (A) WIDs from irradiated 3^rd^ instar WT larvae were immunostained to visualize the dying apoptotic cells (cDcp-1; red; active effector caspases). Shown are representative images of the WIDs displaying an increase in the number of dying epithelial cells already at 4 hpi, which significantly rises by 24 hpi and is dramatically reduced by 48 hpi due to clearance of the apoptotic corpses. Note that 20 Gy of X-rays were used for irradiation throughout this study. (B) Quantification of the number of dying cells in WIDs represented in (A) is presented as the area occupied by dying cells divided by the total WID area, multiplied by 100 to express the percentage of dying cells. (C) The graph depicts total size of the WIDs shown in (A) and presented as the total area in square micrometers (µm²). Note the restoration to normal size at 48 hpi compared to non-irradiated (Non-irrad.) WID counterparts. (D and E) Schematic representations of the Dronc activity genetic reporters *_R_DBS* and *_D_DBS* are shown before (upper) and after (lower) cleavage by Dronc, along with the consequent nuclear translocation of either Histone-GFP (D) or the QF transcription factor (E). The location of the Drice catalytic cysteine at position 211, which was substituted by alanine, is indicated, along with the site of cleavage by Dronc (TETD’G). In the nucleus, the QF transcription factor drives the expression of Gal4, which in turn activates the expression of a gene of interest, such as the nuclear red fluorescent protein (nRFP) RedStinger (E). (F) Irradiated WIDs expressing the *_R_DBS* reporter were immunostained to visualize the dying apoptotic cells (cDcp-1; red). The images show large areas of WIDs with exclusive expression of green fluorescent chromatin (nuclear Histone-GFP) in dying cells at both 4 and 24 hpi. The outlined areas (white squares) are magnified in the corresponding insets. (G) Irradiated WIDs expressing both the *_R_DBS* and *_D_DBS* reporters, along with a *QUAS-Gal4* adapter and a *UAS-RedStinger*, are shown. The images depict large areas of WIDs displaying cells that have activated both reporters, as well as with only one of the reporters activated, at 4 and 24 hpi. The outlined areas (white squares) are magnified in the corresponding insets, which display examples of cells activating one reporter (green chromatin; *_R_DBS*), the other (nuclear red; *_D_DBS*), or both reporters. For the graphs in (B and C), *p* values were calculated using an unpaired Student’s t-test, two-tailed distribution. All data points, including outliers, are presented in box plot format. The minimum value is represented by the lower whisker bound, and the maximum value is represented by the upper whisker bound. The center line of the box denotes the median, with the lower and upper box bounds representing the medians of the lower and upper halves of the dataset, respectively. Each dot corresponds to a single WID measurement, reflecting the number of examined biologically independent WIDs (n). \**p* < 0.05; \*\**p* < 0.01; \*\*\*\**p* < 0.0001; ns (non-significant).

We next sought a specific marker to label the apoptosis-resistant cells that drive the regeneration of the WID. To achieve this, we examined a genetic reporter sensor for the activity of the initiator caspase Dronc. Beyond its role in initiating apoptosis by cleaving and activating effector caspases, Dronc has also been reported to mediate non-apoptotic cellular processes, including AiP. ^38,63,65,75,76^ We hypothesized that a delayed-reporting sensor of Dronc activity, which activates later after exposure to ionizing radiation (IR), might more specifically label surviving cells. By contrast, early-reporting sensors primarily detect apoptotic cells and may overshadow the detection of surviving cells. To investigate this hypothesis, we examined two related *Drice-based sensors* of Dronc activity: a rapid reporter sensor (*_R_DBS*) and a delayed reporter sensor (*_D_DBS*). Originally described in, ^77^ these genetic sensors consist of two key components: a common N-terminal Dronc recognition element and two distinct C-terminal reporting elements (Figures 1D and 1E). The Dronc recognition element features a mouse CD8 transmembrane domain fused with a truncated, inactive version of the effector caspase-3 ortholog Drice. The reporting elements consist of either a Histone H2B fused to GFP (Histone-GFP; in the *_R_DBS*) or a QF transcription factor from the binary Q-system (*_D_DBS*). Upon cleavage by Dronc, the reporting elements translocate from the cell membrane to the nucleus. In cells expressing the *_R_DBS*, this movement is quickly detectable due to the chromatin being fluorescently labeled with GFP. In contrast, in our setup, detecting *_D_DBS* sensor cleavage requires the presence of *QUAS-Gal4* and *UAS*-dependent fluorescent protein constructs. This setup produces a fluorescent signal only after two rounds of transcription and translation - first to produce Gal4 and then the fluorescent protein [such as the nuclear red fluorescent protein (nRFP) RedStinger]. Since most cellular stresses result in the inhibition of cap-dependent translation, ^78–81^ we anticipated that the *_D_DBS* sensor setup would facilitate the delayed detection of Dronc activity. Of note, cleavage of the *_D_DBS* sensor could also be detected following a single round of transcription and translation when used with a *QUAS-*dependent fluorescent protein construct.

To explore the *_D_DBS* reporter, we first examined WIDs from irradiated larvae (20 Gy X-rays) that ubiquitously expressed the early reporter *_R_DBS*. In line with the idea that a rapid sensor of Dronc activity marks apoptotic cells, Histone-GFP translocation to the chromatin was exclusively observed in apoptotic cells at both 4 and 24 hpi, as confirmed by immunostaining with the anti-cleaved Dcp-1 antibody (Figure 1F). Interestingly, irradiated WIDs expressing both *_R_DBS* and *_D_DBS*, along with the *QUAS-Gal4* adapter and *UAS-RedStinger* transgenic constructs, exhibited numerous dying cells with GFP-labeled chromatin (indicating cleaved *_R_DBS*) but no expression of nRFP (indicating cleaved *_D_DBS*) at both 4 and 24 hpi (Figure 1G). In contrast, far fewer nRFP-expressing cells were detected at both time points. However, both the overall number of nRFP-positive cells and the number of nRFP-positive cells with GFP-negative chromatin increased at 24 hpi compared to the 4 hpi time point (Figure 1G). Taken together, these results suggest that, following irradiation of the WID, some epithelial cells activate Dronc and persist for at least up to 24 hpi.

We next asked whether the cleaved *_D_DBS*-positive cells are apoptosis-resistant surviving cells or if they are cells with a delayed but ultimately inevitable demise. To address this question, we similarly immunostained irradiated WIDs expressing the *_D_DBS* reporter along with *QUAS-Gal4* and *UAS-Venus* transgenic constructs (Figure 2A) to identify apoptotic cells marked by cDcp-1, as well as cleaved *_D_DBS*-positive cells labeled with the yellow fluorescent protein Venus. In non-irradiated WIDs, we observed only a few cDcp-1-positive dying cells, as expected, and only a few Venus-positive small cell clones, with the two populations being mutually exclusive (Figure 2Bi). The observation that Venus-positive cells appeared as small clones suggests that, rather than undergoing apoptosis, some cells activate Dronc and still undergo a few rounds of division. At 4 hpi, we observed a dramatic increase in the number of apoptotic cells, but only a modest rise in the number of cleaved *_D_DBS*-positive cells and clones (Figures 2Bii, v, vi). At 24 hpi, there was a significant increase in the number of cleaved *_D_DBS*-positive cells and clones, most of which showed no cDcp-1 expression (Figures 2Biii, vii, viii). This suggests that the *_D_DBS* is more effective at detecting surviving cells that have activated Dronc, rather than the dying cells. Furthermore, by 48 hpi, both the number and size of *_D_DBS*-positive cell clones had increased, with most of these clones being cDcp-1-negative (Figure 2Biv). This indicates that not all irradiated cells that activate Dronc undergo apoptosis; some of these cells survive, divide, and persist throughout the WID even two days after irradiation. Finally, this phenomenon is not limited to WIDs; *_D_DBS*-positive cell clones were also observed in irradiated halteres, legs, and eye-antenna imaginal discs at 48 hpi (Figure 2C). Notably, in the eye-antenna imaginal disc, this phenomenon was largely confined to the antennal region, consistent with the fact that the eye region is already undergoing photoreceptor specification and differentiation at that stage.^82^ Collectively, these results reveal a natural population of <u>D</u>ronc-activating <u>a</u>poptosis resistant <u>e</u>pithelial (DARE) cells scattered throughout imaginal disc tissues. These cells increase in number 24 hours after irradiation and are identified using a delayed genetic reporter of Dronc activity.

**Figure 2.**
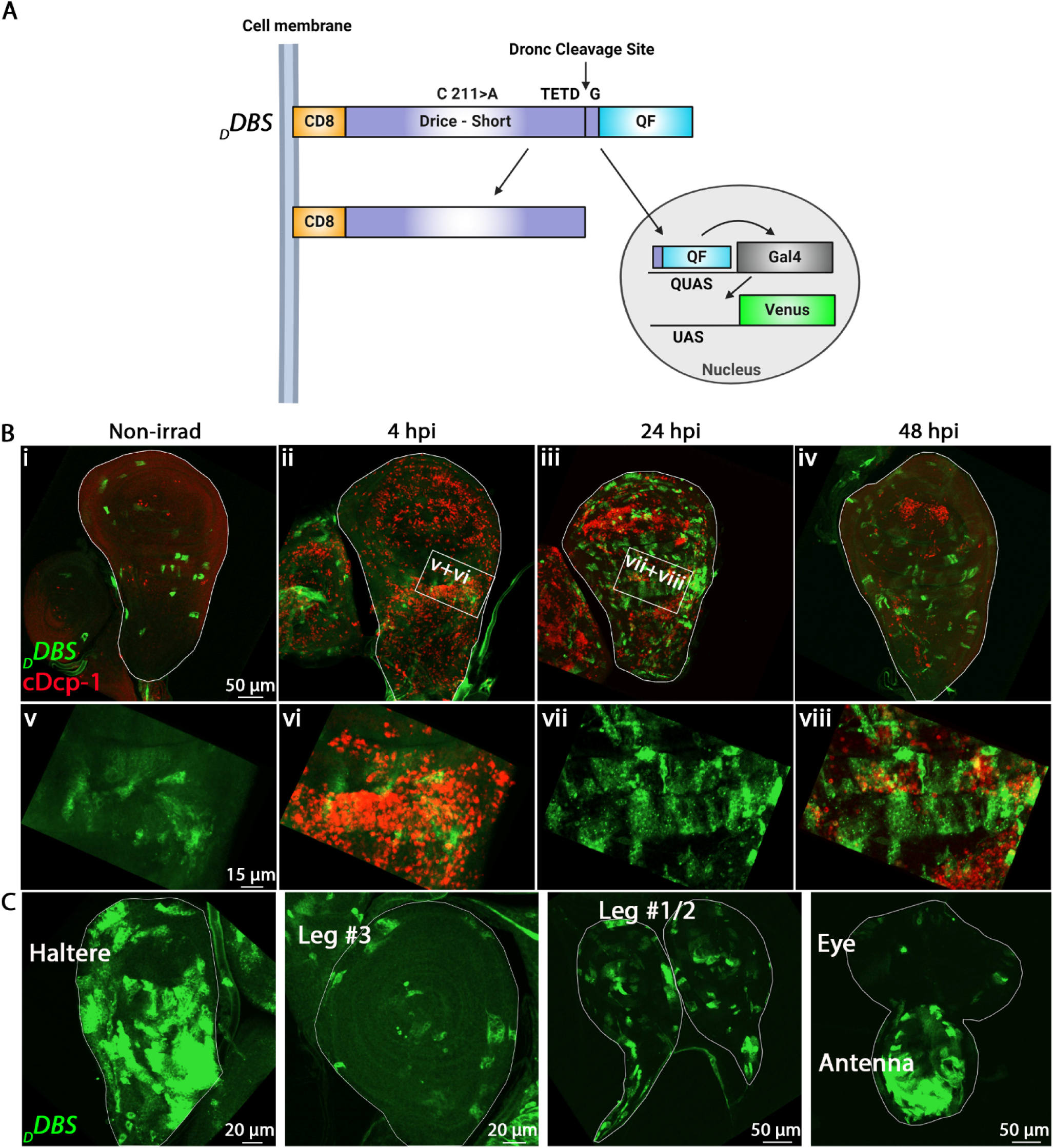
The *_D_DBS* reporter primarily labels apoptosis-resistant cells surviving even 48 hpi. (A) Schematic representation of the *_D_DBS* reporter combined with a *UAS-Venus* transgene. (B) Irradiated WIDs expressing the transgenic combination described in (A) were immunostained to visualize the dying apoptotic cells (cDcp-1; red). The images show representative WIDs before and at the indicated time points after irradiation, displaying largely mutually exclusive expressions of active *_D_DBS* (Venus; green) and cDcp-1 at 4 and 24 hpi, the periods of massive cell death. Note the increase in Venus-positive cells at 24 and 48 hpi. The outlined areas (white rectangles) are magnified in the row below. (C) Representative images of 48 hpi imaginal discs of the indicated types, expressing the transgenic combination described in (A). Cells that activated the reporter (Venus; green) are readily detected, occupying large areas of the tissues.

### DARE Cells Extensively Proliferate between 24-48 hpi and Regenerate the WID

The observed increase in DARE cells at 24 hpi, along with a significant rise in the number and size of fluorescently dimmer DARE cell-derived clones at 48 hpi, suggests that DARE cells may first appear around 24 hpi due to delayed Dronc activation and subsequently proliferate from 24-48 hpi, causing dilution of the fluorescent protein with each cell division. To test this possibility, we employed a genetic system called G-TRACE to track the lineages of dividing cells while avoiding the dilution of the fluorescent protein in the resulting daughter cells (Figure 3A). ^83^ The G-TRACE system consists of three transgenic constructs: a *UAS-nRFP* for real-time red fluorescence, a *UAS-FLP* for Flippase expression, and a *FRT-stop-FRT-nEGFP* cassette for cell lineage tracing with green fluorescence. Combining the *_D_DBS* reporter with the G-TRACE system means that DARE cells should exhibit intense red or yellow (red plus green) fluorescence. In contrast, as DARE cells divide, their daughter cells should maintain constant green fluorescence intensity while showing a gradual decrease in red fluorescence with each cell division (Figure 3A).

**Figure 3.**
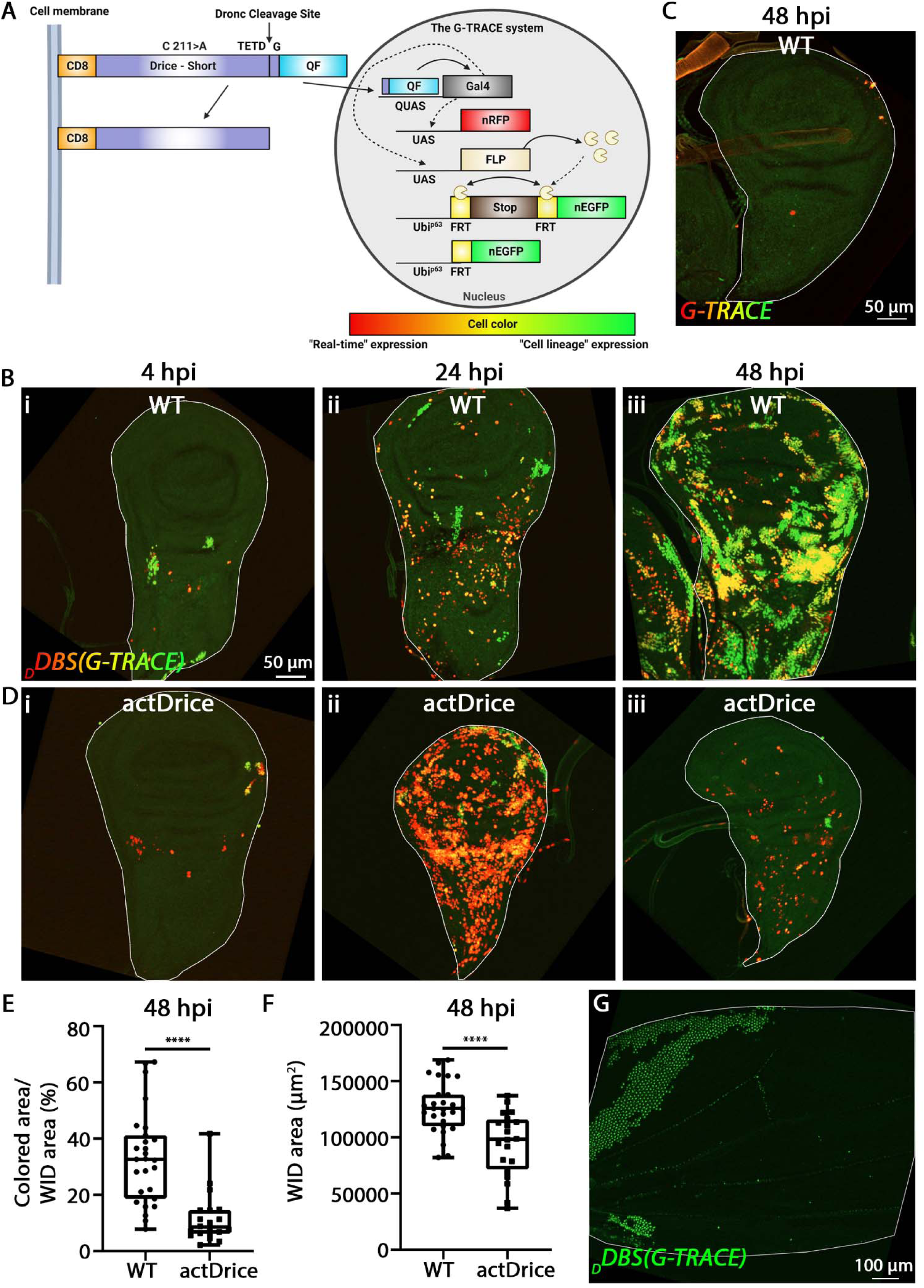
Multiple DARE cells appear at 24 hpi and extensively proliferate over the following 24 hours to regenerate the WID. (A) Schematic representation of the *_D_DBS* reporter combined with the G-TRACE genetic system for cell lineage tracking. Early DARE cells are expected to exhibit red fluorescence (real-time expression) or yellow fluorescence (red combined with the initiation of cell lineage green expression). Large DARE cell clones should display only green fluorescence, as the red fluorescence gradually diminishes with each round of cell division. (B) Irradiated WIDs expressing the transgenic combination described in (A) and monitored at 4, 24, and 48 hpi. Note that newly appearing DARE cells increase at 24 hpi and continue to appear at more advanced time points, as indicated by the presence of yellow cells at 48 hpi. (C) A 48 hpi WID expressing only the G-TRACE system is shown to control for possible system leakage. The absence of fluorescent cells in this WID indicates that the fluorescence of DARE cells in (B) is exclusively attributed to the activation of the *_D_DBS* reporter. (D) Irradiated WIDs, as in (B), expressing a *UAS-actDrice* transgenic construct in the DARE cells to produce a constitutively active Drice (actDrice) that triggers apoptosis and specific ablation of these cells. See Figures S1A-S1C for more details about the actDrice structure and its efficiency. Note the accumulation of dying DARE cells at 24 hpi (red cells with essentially no or minimal green expression) and their clearance from the tissue by 48 hpi, by which time most apoptotic cells in the WID are also cleared (see Figure 1A). (E) Quantification of the number of DARE cells and their derivative daughter cells in 48 hpi WIDs, as represented in (B and D), is shown. This is presented as the area occupied by fluorescent cells (red, green, or both) divided by the total WID area, multiplied by 100 to express the percentage of the WID area occupied by DARE cells and their derived clones. (F) The graph depicts the total size of the 48 hpi WIDs shown in (B and D), presented as the total area in square micrometers (µm²). Note the significant decrease in WID size following DARE cell ablation, indicating the crucial role of DARE cells in compensatory proliferation-mediated tissue regeneration. (G) The image depicts an early eclosed adult fly wing corresponding to the irradiated larvae described in (B). Residual DARE cell clones are still detectable in the wing (green) just before their normal removal, along with the entire wing epithelium, by apoptosis. For the graphs in (E and F), *p* values were calculated using an unpaired Student’s t-test, two-tailed distribution. The graphs were generated and presented as in Figure 1B. \*\*\*\**p* < 0.0001.

Consistent with our observation that tissue regeneration in irradiated WIDs predominantly occurs between ∼24-48 hpi (Figure 1C), only a few red and yellow cells, along with small green cell clones, are detected at 4 hpi (Figure 3Bi). By 24 hpi, there is a significant increase in the number of red and yellow cells, which are dispersed throughout the WID (Figure 3Bii). Critically, the number and size of yellow/green cell clones increase dramatically at 48 hpi, covering approximately 40-50% of the WID area, suggesting extensive proliferation of DARE cells between 24-48 hpi (Figures 3Biii and 3E). Note that the fluorescent cells are exclusively DARE cells and derivative clones, as no fluorescent cells are detected at 48 hpi in irradiated WIDs lacking the *_D_DBS* reporter, indicating that there is no leakiness from the G-TRACE system (Figure 3C).

The observation that DARE cell-derived clones occupy a large portion of the regenerated WID at 48 hpi, suggests an important role of the DARE cells in tissue regeneration. However, the regenerated WID also contains <u>n</u>on-Dronc-activating <u>a</u>poptosis resistant <u>e</u>pithelial (NARE) cells (indicated by ‘dark’ areas in Figure 3Biii). We therefore investigated whether the DARE cells are crucial for WID regeneration or if NARE cells can promote normal regeneration in the absence of DARE cells. We ablated the DARE cells using a constitutively active Drice transgene (*actDrice*) that strongly induces apoptosis when ectopically expressed (Figures S1A-S1C). Irradiated WIDs expressing the *_D_DBS*/G-TRACE systems along with a *UAS-actDrice* transgenic construct contained only a few predominantly red-labeled DARE cells and a few rare small clones at 4 hpi, indicating that this combined system effectively triggers apoptosis in the DARE cells (Figure 3Di). Consistent with this, numerous, almost exclusively red-labeled DARE cells and a few rare clones were detected at 24 hpi (Figure 3Dii). By 48 hpi, these cells were largely cleared from the tissue, which showed no DARE cell-derived clones (Figures 3Diii and 3E). Importantly, the ablation of the DARE cells led to a significant reduction in WID area at 48 hpi, indicating that NARE cells cannot compensate for the loss of DARE cells and underscoring the critical role of DARE cells in tissue regeneration (Figure 3F).

To directly visualize mitotic cells in the regenerating tissue, irradiated WIDs expressing the *_D_DBS*/G-TRACE systems were immunostained with an antibody specific for Phosphohistone H3 (pHH3), a marker of mitosis. ^84^ Monitoring WIDs at 24 hpi, when multiple DARE cells along with DARE cell-derived clones become apparent, revealed multiple mitotic cells distributed relatively uniformly throughout the tissue (Figure S1E). However, closer examination of the dividing cells showed that many were located not within the cell clone masses but rather at their periphery (Figure S1F). Furthermore, consistent with the idea that DARE cells are crucial for NARE cell proliferation, these pHH3-positive cells often lacked fluorescent signals, indicating that they are NARE cells potentially receiving mitogenic signals from adjacent DARE cells (Figure S1G). Taken together, we conclude that following irradiation, some of the DARE cells at 24 hpi are induced to extensively proliferate and regenerate the WID.

### DARE Cell Clones Support Normal Adult Wing Development and Resist Apoptosis Induced by Re-Irradiation

Shortly after eclosion, the wing epithelium of young adult flies is typically removed by apoptosis. ^85^ To investigate whether DARE cell clones contribute to the formation of normal adult wings, we monitored the flies immediately after eclosion. Indeed, following larval-stage irradiation, some DARE cell-derived green clones could still be detected shortly after eclosion in the fly wings expressing the *_D_DBS*/G-TRACE systems, indicating that these clones contribute to functional wing development (Figure 3G).

The finding that DARE cells survive irradiation, which induces apoptosis in many similar epithelial cells in the WID, raises the question of whether this resistance is specific to the early DARE cells or if it is also inherited by their daughter cells. To address this question, irradiated WID carrying the *_D_DBS*/G-TRACE systems were re-irradiated at 48 hpi. The tissues were examined 4 hours after the second irradiation, following immunostaining with anti-cDcp-1 to identify apoptotic cells. Interestingly, while dying epithelial cells cover approximately 25% of the total WID area 4 hours after the first irradiation (Figure 1A and 1B), this percentage decreases by about half to roughly 13% 4 hours after the second irradiation (Figures S2A and S2E). Moreover, detailed analysis of cDcp-1 expression in DARE cells and their daughter cells revealed that the likelihood of these cells dying after re-irradiation is approximately 5%, compared to the expected 25% if cell death were random (Figures S2B-S2D and S2F). We therefore conclude that the resistance to apoptosis observed in DARE cells is inherited by their daughter cells, suggesting the presence of a molecular memory.

### Dronc, but Not Dark or Effector Caspases, Is Essential for DARE Cell Proliferation and WID Regeneration

The detection of DARE cells due to delayed Dronc activity raises the critical question of whether Dronc is necessary for DARE cell proliferation and tissue regeneration. To test this possibility, we used a specific RNAi transgene (Figure S1D) to knock down *dronc* in DARE cells of irradiated WIDs expressing the *_D_DBS*/G-TRACE systems. Downregulation of Dronc in DARE cells significantly reduced their proliferation between 24-48 hpi, resulting in much smaller DARE cell-derived clones that occupy less WID area at 48 hpi compared to WT counterparts (Figure 4A-4C). Importantly, the reduced proliferation of DARE cells following *dronc* knockdown was not compensated by increased proliferation of NARE cells, resulting in a significant reduction in WID size (Figure 4D). Given that Dronc activity is required for the initial cleavage of the *_D_DBS* reporter and that the *dronc^Ri^*is expressed in DARE cells only after this cleavage, the observed retardation in DARE cell proliferation likely even underestimates the true requirement of Dronc for DARE cell proliferation and tissue regeneration. Collectively, we conclude that Dronc activity is essential for the proliferation of DARE cells.

**Figure 4.**
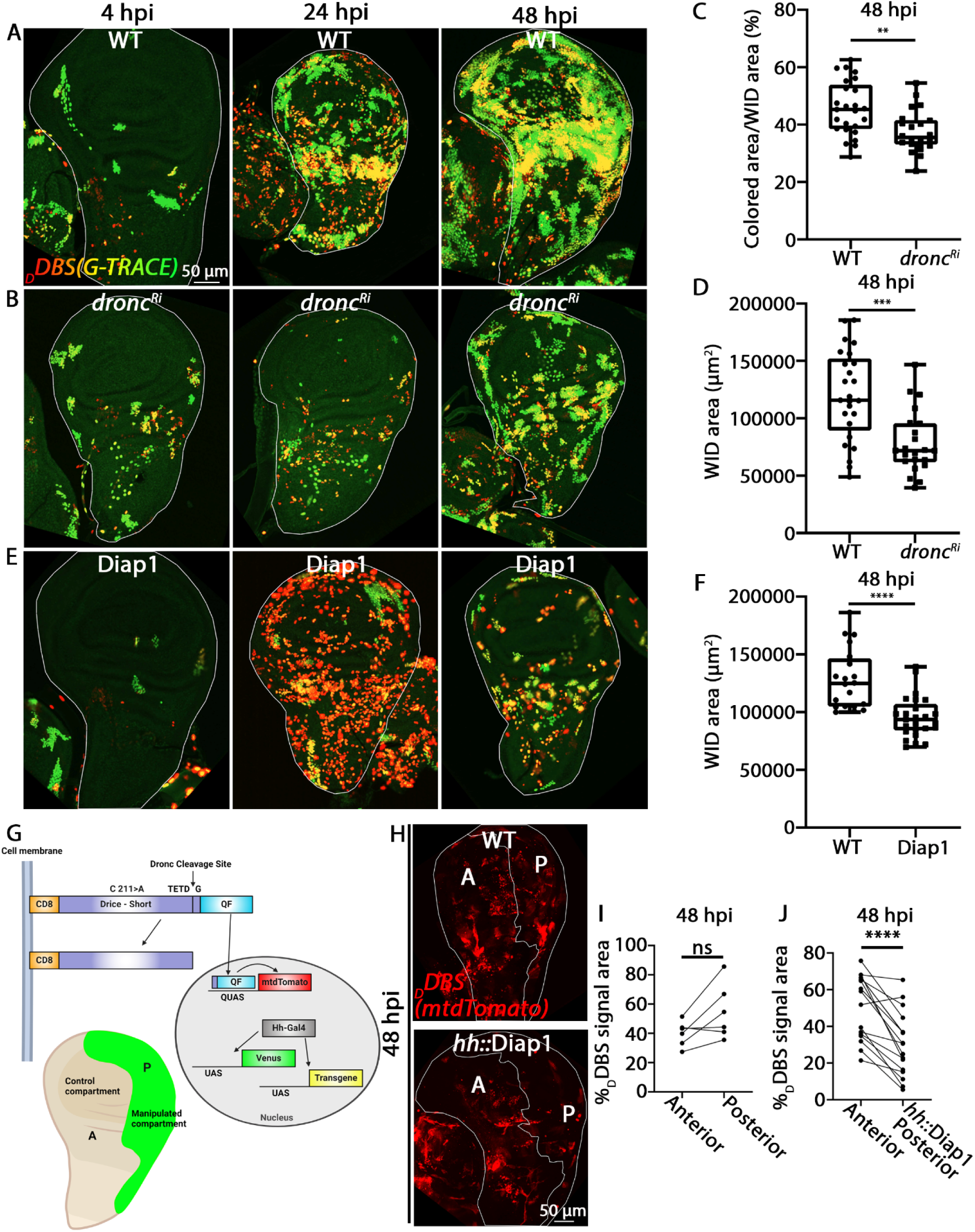
Dronc is required for DARE cell proliferation and tissue regeneration. (A and B) Irradiated WIDs expressing the transgenic combination described in Figure 3A, either without (A) or with a *UAS-dronc^Ri^* transgene (B), were monitored at 4, 24, and 48 hpi. Note the decrease in the number of dividing DARE cells (green) at both 24 and 48 hpi when *dronc* is targeted by a specific RNAi transgene. (C) Quantification of the number of DARE cells and their derivative daughter cells in 48 hpi WIDs, as represented in (A and B), is shown and presented as in Figure 3E. (D) The graph depicts the total size of the 48 hpi WIDs shown in (A and B), presented as the total area in square micrometers (µm²). Note the significant decrease in WID size following downregulation of Dronc, indicating an essential role of Dronc in DARE cells for tissue regeneration. (E) Irradiated WIDs of the genotype described in (A) that also contain a *UAS-diap1* transgene, which overexpresses the Dronc and effector caspase activity inhibitor in DARE cells. Note that, while Diap1 blocks the natural apoptosis of many DARE cells through its inhibitory effect on effector caspase activity, leading to their accumulation at 24 hpi (red cells), it also causes a dramatic reduction in the size and number of DARE cell clones (green) at 48 hpi due to its inhibition of Dronc activity. See also Figure S3B for expression of P35, which specifically inhibits the effector caspases but not Dronc. (F) The graph depicts the total size of the 48 hpi WIDs shown in (E) compared to control WIDs of the genotype described in (A), presented as the total area in square micrometers (µm²). Note the significant decrease in WID size, reflecting the tissue’s impaired ability to regenerate. Since overexpression of P35 in the DARE cells Figure S3D does not affect tissue regeneration, the effect caused by Diap1 overexpression is attributed to its potent inhibition of Dronc activity. (G) Schematic representation of the dual binary system, featuring the *_D_DBS* reporter combined with a *QUAS-mtdTomato* transgene for detection of DARE cells (red), and an *hh-Gal4* transgenic construct to drive expression of a transgene of interest, such as a fluorescent protein or *diap1*, exclusively in the posterior compartment of the WID. (H) Irradiated WIDs of the genotype described in (G) either without (top) or with the *hh-Gal4* and *UAS-diap1* transgenes (bottom) at 48 hpi, reveal that Diap1 overexpression leads to a dramatic decrease in the number and size of DARE cell clones in the posterior compartment compared to the anterior compartment of the WID. (I and J) Before-and-after presentation of individual values shown as the percentage of area occupied by DARE cell clones in the posterior and anterior compartments of the WIDs represented in (H). Each pair of connected dots represents measurements from a single WID, with each dot indicating the data for either the anterior or posterior compartments. The DARE cell-occupied areas are presented as the areas covered by red fluorescent cells in each compartment, divided by the total compartment area, and multiplied by 100. Note the natural trend (albeit non-significant) of DARE cells occupying larger areas in the posterior compartment compared to the anterior compartment (I), and the significant reversal of this trend when Diap1 is overexpressed in the posterior compartment (J). For the graphs in (C, D, and F), *p* values were calculated using an unpaired Student’s t-test, two-tailed distribution. The graphs were generated and presented as in Figure 1B. \*\**p* < 0.01; \*\*\**p* < 0.001; \*\*\*\**p* < 0.0001. For the graphs in (I and J), *p* values were calculated using a paired Student’s t-test, two-tailed distribution. All data points, including outliers, are shown in a plot displaying individual values. \*\*\*\**p* < 0.0001; ns (non-significant).

An alternative way to attenuate Dronc activity in the DARE cells is by expressing the *Drosophila* caspase inhibitor Diap1, which binds to and inhibits both Dronc and the effector caspases. ^22,38–40^ Consistent with reduced Dronc activity, significant attenuation in DARE cell proliferation was observed, evidenced by the near absence of DARE cell-derived clones at 24 hpi and the presence of only a few small clones at 48 hpi in irradiated WIDs expressing a *UAS-diap1* transgenic construct (Diap1(1) in Figure S1D) along with the *_D_DBS*/G-TRACE systems (Figure 4E). Interestingly, multiple, red-labeled DARE cells were detected at 24 hpi with only a few yellow cells and nearly complete absence of green cell clones following Diap1 expression (Figure 4E). This is likely due to the potent inhibition of Dronc, which strongly attenuates DARE cell proliferation and clone formation, as well as Diap1’s role as a potent apoptosis inhibitor, which prevents the natural death of some DARE cells by simultaneously inhibiting Dronc and the effector caspases. Finally, similar to the effect of Dronc downregulation, DARE cells expressing Diap1 failed to effectively regenerate the irradiated WID, as evidenced by the significant reduction in WID size at 48 hpi (Figure 4F).

Although inactivation of Dronc in DARE cells using specific RNAi and Diap1 overexpression led to a significant reduction in the number and size of DARE cell-derived clones at 48 hpi, clone generation was not completely blocked. This is presumably because transgene expression in DARE cells occurs only after the initial cleavage of the *_D_DBS* reporter by Dronc, as well as natural variations in transgene expression efficiency and function among individual cells. To evaluate the contribution of the natural variation factor to the incomplete block of DARE cell clone formation, we overexpressed Diap1 (Diap1 (2) in Figure S1D) in the entire posterior region of the WID using the *hedgehog* (*hh*)-*Gal4* driver. DARE cells were visualized at 48 hpi using a *QUAS-mtdTomato* transgenic construct, which expresses the membrane-targeted red fluorescent protein tdTomato following cleavage of the *_D_DBS* reporter by Dronc and the subsequent nuclear translocation of the QF transcription factor (Figure 4G). Irradiated WIDs with this transgenic combination showed a significant decrease in the number and size of DARE cell-derived clones in the posterior region compared to the unmanipulated anterior region of the WID following Diap1 overexpression (Figures 4H-4J). Nevertheless, despite the dramatic effect - unobserved in a similar control background - some small cell clones were still detected in the posterior region. These clones may result from natural variations in Diap1 overexpression and efficiency among individual DARE cells. Additionally, they could have originated from clones generated near the anterior/posterior border, where they might still receive mitogenic signals from unmanipulated DARE cells in the anterior region.

Since Dronc activates the effector caspases and Diap1 also inhibits them, we wondered whether the strong inhibitory effect of Diap1 overexpression on DARE cell-derived clone formation might be, at least in part, due to a requirement for the effector caspases in promoting DARE cell proliferation. To investigate this possibility, we overexpressed the baculovirus P35 protein in DARE cells using the *_D_DBS*/G-TRACE systems. P35 can potently inhibit apoptosis by binding to and inhibiting both Drice and Dcp-1, but not Dronc (Figure S1D). ^29,36,39,86–88^ However, P35 overexpression in DARE cells did not affect cell proliferation or overall WID regeneration, as indicated by normal clone formation and WID size at 48 hpi (Figures S3A, S3B and S3D). Notably, although P35 overexpression did not affect clone formation, it did inhibit the natural death of some DARE cells due to its potent apoptosis-inhibiting properties, as evident by the accumulation of multiple individual red/yellow-labeled non-dividing DARE cells at 24 and 48 hpi (Figure S3B). We can conclude that effector caspases in DARE cells do not influence their proliferation or tissue regeneration. This suggests that the impact of Diap1 on DARE cell proliferation is entirely attributable to the inhibition of Dronc.

As a key activator of Dronc during apoptosis, the apoptosome Apaf-1 adapter ortholog Dark is essential for nearly all apoptotic events in *Drosophila*. ^31,32,34,35,37,89^ To determine if non-lethal activation of Dronc in DARE cells also requires Dark, we expressed a *dark* RNAi transgene (Figure S1D) in the DARE cells using the *_D_DBS*/G-TRACE systems and monitored its effects on cell proliferation and WID regeneration. Unexpectedly, *dark* knockdown had no significant effect on DARE cell proliferation or WID regeneration at any of the examined time points after irradiation (Figures S3A and S3C-S3E). Therefore, non-lethal Dronc activation in DARE cells likely occurs independently of the Dark apoptosome.

### Myo1D, an Unconventional Myosin Interacting with Dronc, Hinders Effector Caspase Activation in DARE Cells

An important question arising from our findings is how DARE cells survive despite activating Dronc. To begin addressing this, we first investigated whether DARE cells exhibit effector caspase activity. To reduce background interference from the many dying cells and improve cellular resolution, we chose to express a genetic reporter specific to effector caspase activity exclusively in DARE cells, instead of using a global immunostaining approach for activated effector caspases. Called *CPV*, this well-established reporter of Drice and Dcp-1 activity consists of three components: an N-terminal mouse CD8 transmembrane domain, a short polypeptide of the human PARP protein containing a specific effector caspase cleavage site, and a C-terminal Venus fluorescent protein. ^29^ Upon apoptosis induction and cleavage by effector caspases, Venus translocates from the cell membrane to the cytoplasm. This translocation can be readily detected through the newly exposed PARP epitope (cPARP) using an anti-cleaved PARP antibody, as demonstrated by the expression of *CPV* in the pouch region of 3 hpi WIDs (Figure 5A and 5B). Monitoring irradiated WIDs expressing the *_D_DBS* reporter, along with *QUAS-Gal4* and *UAS*-*CPV* transgenic constructs, and immunostained for cPARP, revealed that most Venus-positive DARE cells exhibit low to no effector caspase activity at 4, 24, and 48 hpi (Figure 5C). While these findings clarify how DARE cells evade apoptotic death, it remains unclear why effector caspases are silent in cells that activate Dronc.

**Figure 5.**
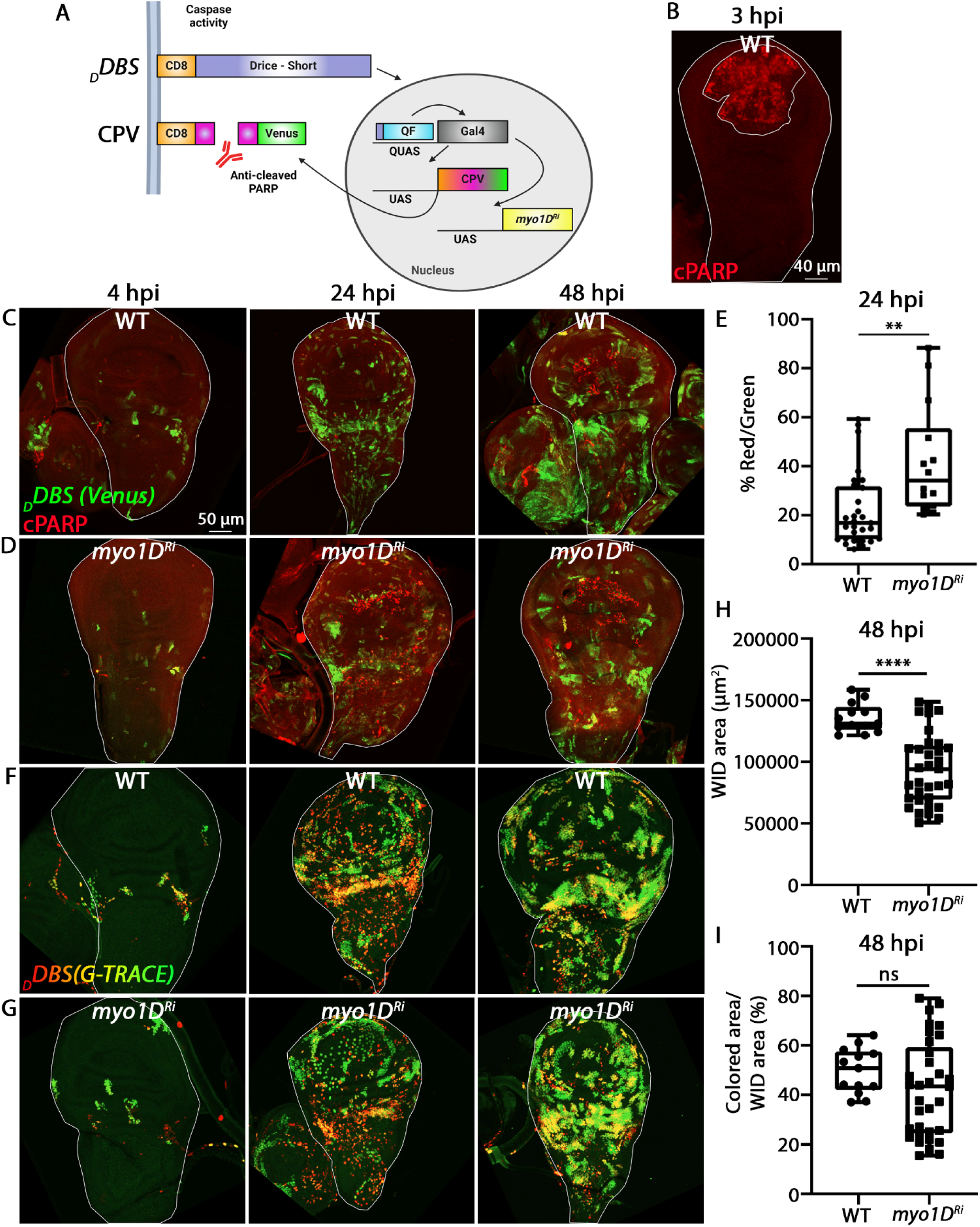
*myo1D* knockdown in DARE cells triggers effector caspase activation and apoptosis. (A) Schematic representation of the *_D_DBS* reporter combined with the *CPV* reporter of effector caspase activity and a *UAS-myo1D^Ri^* transgene. Gal4 expression, following cleavage of the *_D_DBS* reporter by Dronc and QF nuclear translocation, induces the expression of the *CPV* reporter and an RNAi against *myo1D* specifically in DARE cells. Cleavage of the CPV reporter by Drice and Dcp-1 after the caspase-3 consensus site DEVD, located within a 40-amino-acid polypeptide of the human PARP protein (indicated in magenta), exposes a new epitope at the C-terminal fragment of the PARP polypeptide. This cleavage releases the Venus yellow fluorescent protein into the cytoplasm, an event that is readily detected using an anti-cleaved human PARP antibody. (B) An example of an irradiated WID expressing the *CPV* reporter in the pouch region (using the *sal-Gal4* driver) and immunostained to reveal cleaved PARP (cPARP; red). (C and D) Irradiated WIDs expressing the transgenic combination described in (A), either without (C) or with the *UAS-myo1D^Ri^* transgene (D), were immunostained for cPARP and monitored at 4, 24, and 48 hpi. DARE cells are visualized by the Venus fluorescence of the *CPV* (green) and/or by cPARP (red) after its cleavage by the effector caspases. Note that cells in advanced apoptotic stages tend to lose Venus fluorescence. (E) Quantification of the number of DARE cells that activated effector caspases (i.e., dying cells; cPARP) in WIDs represented in (C and D). The graph represents the area occupied by cPARP-expressing DARE cells (red) divided by the area covered by DARE cells without effector caspase activity (green), multiplied by 100 to express the percentage of DARE cells exhibiting effector caspase activity. (F and G) Irradiated WIDs expressing the transgenic combination described in Figure 3A, either without (F) or with a *UAS-myo1D^Ri^* transgene (G), were monitored at 4, 24, and 48 hpi. (H) The graph depicts the total size of the 48 hpi WIDs shown in (F and G), presented as the total area in square micrometers (µm²). Note the significant decrease in WID size following downregulation of Myo1D, consistent with the role of this unconventional myosin in protecting DARE cells from excessive effector caspase activation by Dronc and subsequent cell death. (I) Quantification of the number of DARE cells and their derivative daughter cells in 48 hpi WIDs, as represented in (F and G), is shown and presented as in Figure 3E. For the graphs in (E, H, and I), *p* values were calculated using an unpaired Student’s t-test, two-tailed distribution. The graphs were generated and presented as in Figure 1B. \*\**p* < 0.01; \*\*\*\**p* < 0.0001; ns (non-significant).

Beyond its interaction with Dark during apoptosis, Dronc has been reported to bind other regulatory proteins involved in non-apoptotic functions. In both an AiP setup in the eye-antenna imaginal disc and in mature enterocytes of the adult posterior midgut, Dronc has been shown to be sequestered to the basal side of the cell membrane by the class I unconventional myosin, Myo1D. ^68,69^ Hypothesizing that Dronc’s binding to Myo1D might inhibit its effective engagement with and activation of effector caspases, we set out to test this idea. We knocked down *myo1D* in DARE cells using an RNAi from the TRiP library and monitored effector caspase activity using the *CPV* reporter. Strikingly, downregulation of *myo1D* significantly increased effector caspase activity at all examined time points (Figure 5D). Furthermore, this effect was particularly pronounced at 24 hpi, when the number of cPARP-positive cells significantly increased (Figures 5D and 5E). It is important to note that the cPARP-positive cells appeared highly condensed compared to the Venus-positive, cPARP-negative cells, indicating that DARE cells which activate effector caspases undergo apoptosis.

To strengthen this intriguing finding and assess the impact of Myo1D downregulation on WID regeneration, we employed a second, independently generated RNAi transgene targeting *myo1D* from the VDRC KK library, in the background of the *_D_DBS*/G-TRACE systems. Importantly, like *dronc* knockdown, downregulation of Myo1D in the DARE cells resulted in a significant reduction in the overall size of the WID at 48 hpi (Figures 5F-5H). Furthermore, the relative WID area occupied by *myo1D* knockdown DARE cell clones at 48 hpi was similar in percentage to that covered by WT counterparts, indicating that there was no overproliferation of NARE cells at the expense of the DARE cells (Figures 5F, 5G and 5I). Taken together, these findings demonstrate that Myo1D plays a crucial role in protecting DARE cells from excessive caspase activation and apoptosis, thereby facilitating their proliferation and contributing to tissue regeneration.

Another Dronc-binding protein, the unconventional non-muscle myosin 7A (MYO7A) ortholog Myo7A/Crinkled, has been reported to promote Dronc activation in both apoptotic and non-apoptotic contexts. ^90^ Similarly, the eukaryotic translation initiation factor 3 subunit M (EIF3M) ortholog, Eif3m/Tango7, is reported to mediate some of Dronc’s non-apoptotic activities at the cell cortex. ^91^ To explore the roles of Crinkled and Tango7 in regulating Dronc activity during compensatory proliferation, we expressed specific RNAi transgenes targeting these genes in DARE cells using the *_D_DBS* reporter system and monitored effector caspase activity via DARE cell expression of the *CPV* reporter (as illustrated in Figure 5A). Interestingly, while knockdown of *tango7* had no effect on the low residual activity of effector caspases in DARE cells at 24 hpi, downregulation of Crinkled resulted in a significant reduction in the already low effector caspase activity (Figure S4). Thus, whereas one unconventional myosin, Myo1D, inhibits effector caspase activation and ensures DARE cell survival, another, Myo7A/Crinkled facilitates it, which may explain why not all DARE cells survive.

### Regulation of DARE Cell Proliferation and Tissue Regeneration by JNK and TNF Receptors

Studies of AiP and other Dronc-mediated paradigms, both apoptotic and non-apoptotic, have shown that the Jun N-terminal kinase (JNK) signaling pathway is activated downstream of Dronc. ^63,70,75,76,92–97^ To investigate possible involvement of the JNK pathway in proliferation of the DARE cells and tissue regeneration, we first knocked down *basket* (*bsk^Ri^*), the sole *Drosophila jnk* gene, in flies expressing the *_D_DBS*/G-TRACE systems (Figure 6A). Whereas no effect on irradiated WIDs was observed at 4 and 24 hpi following *jnk* downregulation, a significant decrease in both the number and size of DARE cell-derived clones was revealed at 48 hpi (Figures 6B, 6C and 6F). Intriguingly, while inactivation of Dronc in DARE cells inhibited the proliferation of both DARE and NARE cells and impaired tissue regeneration, inhibition of DARE cell proliferation upon *jnk* knockdown was compensated by an increase in NARE cell proliferation, as was reflected in the normal size of WIDs at 48 hpi (Figures 6B, 6C and 6G).

**Figure 6.**
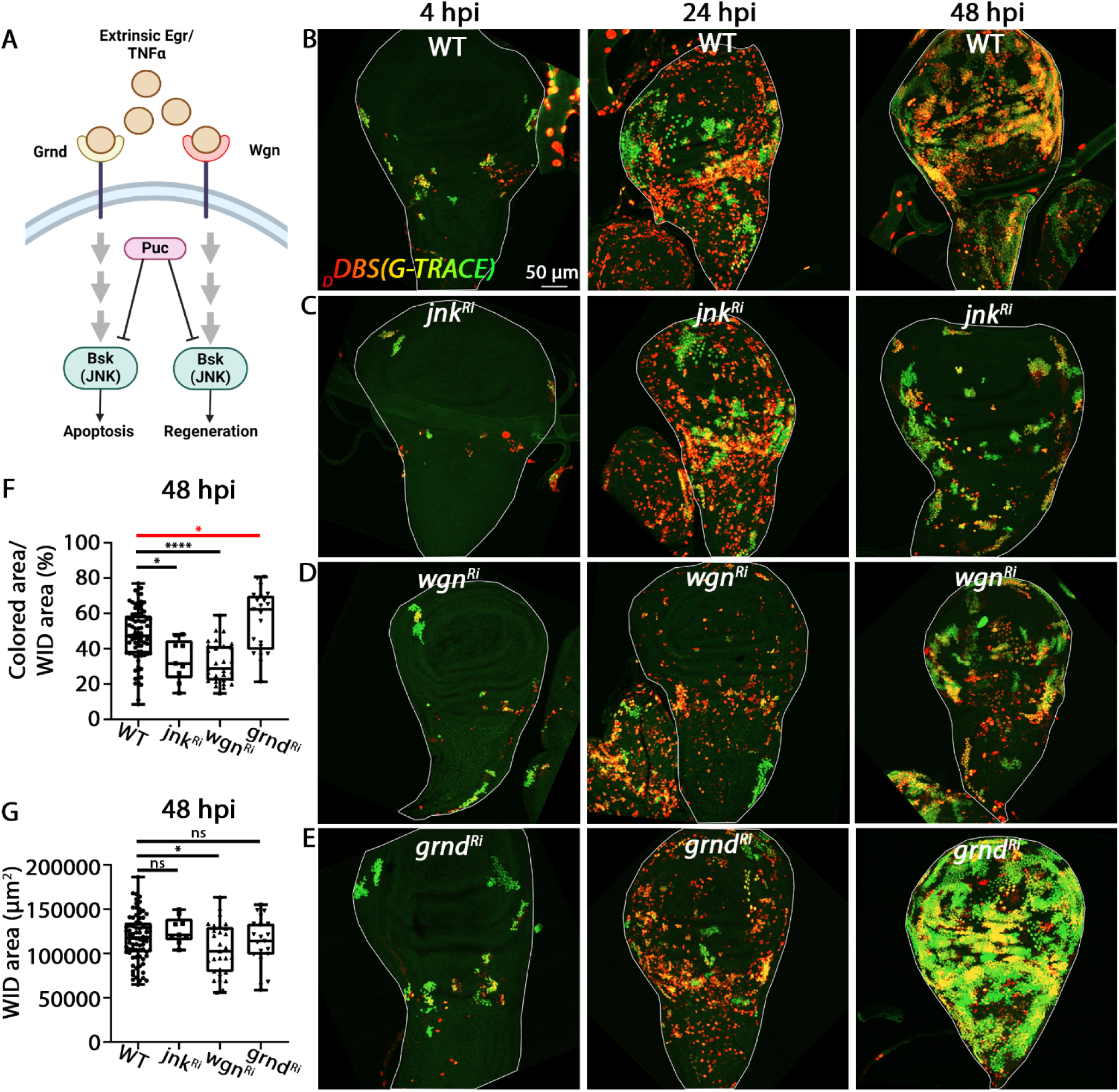
The TNFR>JNK Signaling Pathway Regulates Proliferation of DARE Cells during Compensatory Proliferation. (A) A schematic summary based on existing literature of the TNFR>JNK signaling pathway, highlighting its roles in apoptosis and tissue regeneration, and illustrating the components examined in this study. (B-E) Irradiated WIDs expressing the transgenic combination described in Figure 3A were monitored at 4, 24, and 48 hpi. The conditions included: a control with no additional transgene (B), a *UAS-jnk^Ri^*transgene (C), a *UAS-wgn^Ri^* transgene (D), and a *UAS-grnd^Ri^*transgene (E). Note that no detectable effects were observed at 4 and 24 hpi in any of the tested conditions. However, by 48 hpi, downregulation of JNK and Wgn in the DARE cells - but not Grnd - resulted in a significant reduction in the area occupied by DARE cell clones in the WID. In contrast, knockdown of *grnd* led to a moderate increase in the overall area covered by DARE cell clones in the WID. (F) Quantification of the number of DARE cells and their derivative daughter cells in 48 hpi WIDs, as represented in (B-E), is shown and presented as in Figure 3E. Statistical significance indicating an increase is represented with a red asterisk, while significance indicating a decrease is shown with black asterisks. (G) The graph depicts the total size of the 48 hpi WIDs shown in (B-E), presented as the total area in square micrometers (µm²). Note that, despite the significant effects of downregulating the TNFR>JNK pathways, the sizes of the WIDs remained normal or were only slightly reduced, suggesting that NARE cells overproliferated at the expense of the mutant DARE cells. For the graphs in (F and G), *p* values were calculated using an unpaired Student’s t-test, two-tailed distribution. The graphs were generated and presented as in Figure 1B. \**p* < 0.05; \*\*\*\**p* < 0.0001; ns (non-significant).

To further validate the role of JNK in DARE cell proliferation, we inactivated JNK signaling in the entire posterior region of irradiated WIDs, as illustrated in Figure 4G. Monitoring DARE cell-derived clones overexpressing either a dominant-negative JNK transgene (*jnk^DN^*) or the JNK inhibitor phosphatase Puckered (Puc; Figure 6A) ^98^ at 48 hpi revealed a significant decrease in both the number and size of these clones, compared to the unmanipulated anterior region (Figure S5). Together, these findings indicate that JNK signaling is crucial for cell-autonomous proliferation in DARE cells but is dispensable for non-cell-autonomous proliferation of NARE cells.

JNK activation in *Drosophila* can be triggered through the activation of two TNF receptors, Wengen (Wgn) and Grindelwald (Grnd). ^99–104^ Interestingly, recent findings suggest that Grnd promotes TNFα-induced apoptosis, while Wgn is required for apoptosis-induced cell survival and WID regeneration.^105^ Additionally, Grnd has been reported to facilitate overgrowth of undead epithelial tissue during AiP, likely through JNK activation. ^95^ We therefore decided to examine the potential involvement of TNF receptors in compensatory proliferation by knocking down *grnd* and *wgn* in the DARE cells. Similar to *jnk* knockdown in DARE cells, no significant effects on irradiated WIDs expressing the *_D_DBS/*G-TRACE systems were observed upon knockdown of either receptor at 4 and 24 hpi (Figures 6B, 6D and 6E). However, at 48 hpi, a significant decrease in both the number and size of DARE cell-derived clones was observed in *wgn* knockdown WIDs, whereas *grnd* knockdown resulted in a moderate increase in the number and size of cell clones (Figures 6B, 6D, 6E and 6F). These results suggest that Grnd and Wgn have opposing roles in regulating DARE cell proliferation, with Grnd acting negatively and Wgn positively. Measuring WID area size at 48 hpi revealed no change with *grnd* knockdown and a mild decrease with *wgn* knockdown (Figure 6G). Therefore, in conjunction with the results from *jnk* knockdown, these findings suggest that reduced proliferation in one cell population may lead to compensatory proliferation in the other population.

### Proliferative Homeostasis between DARE and NARE Cells Ensures Balanced Compensatory Growth of the Tissue

Our findings indicate that Dronc activity in DARE cells is essential for the proliferation of both DARE and NARE cells. Additionally, the TNFR>JNK signaling pathway within DARE cells is involved in maintaining a balanced proliferative equilibrium between these two cell populations. However, it remains unclear whether this equilibrium is directly regulated by signals originating from DARE cells downstream of the TNFR>JNK signaling pathway, or if it is an indirect result of regulating the rate of DARE cell proliferation. To address this question, we aimed to attenuate DARE cell proliferation downstream of the TNFR>JNK signaling by overexpressing the *Drosophila* Cyclin E/Cdk2 inhibitor protein, Dacapo (Dap). ^106,107^ Whereas monitoring irradiated WIDs expressing the *_D_DBS* reporter, the *QUAS-Gal4* adapter, a *UAS*-*Dap-GFP* transgenic construct, and a *UAS-RedStinger* for DARE cell detection showed no significant effects at 4 and 24 hpi, a significant reduction in both the number and size of DARE cell clones became evident at 48 hpi (Figures 7A, 7B and 7D). Furthermore, we believe that the effect of Dap overexpression is even more pronounced, as the sparse and small clones observed at 48 hpi may be attributed to inefficient Dap expression in the originating DARE cells, as indicated by the absence of GFP fluorescence in these clones (Figure 7C). Importantly, despite the reduced proliferation of DARE cells, the Dap-overexpressing WIDs maintained an overall normal size at 48 hpi, suggesting that NARE cells were overproliferating at the expense of the DARE cells (Figure 7E). Given that DARE cells overexpressing Dap retain active JNK signaling, these findings suggest that the TNFR>JNK pathway may play an indirect role in regulating the proliferative homeostasis between DARE and NARE cells by modulating the rate of DARE cell proliferation.

**Figure 7.**
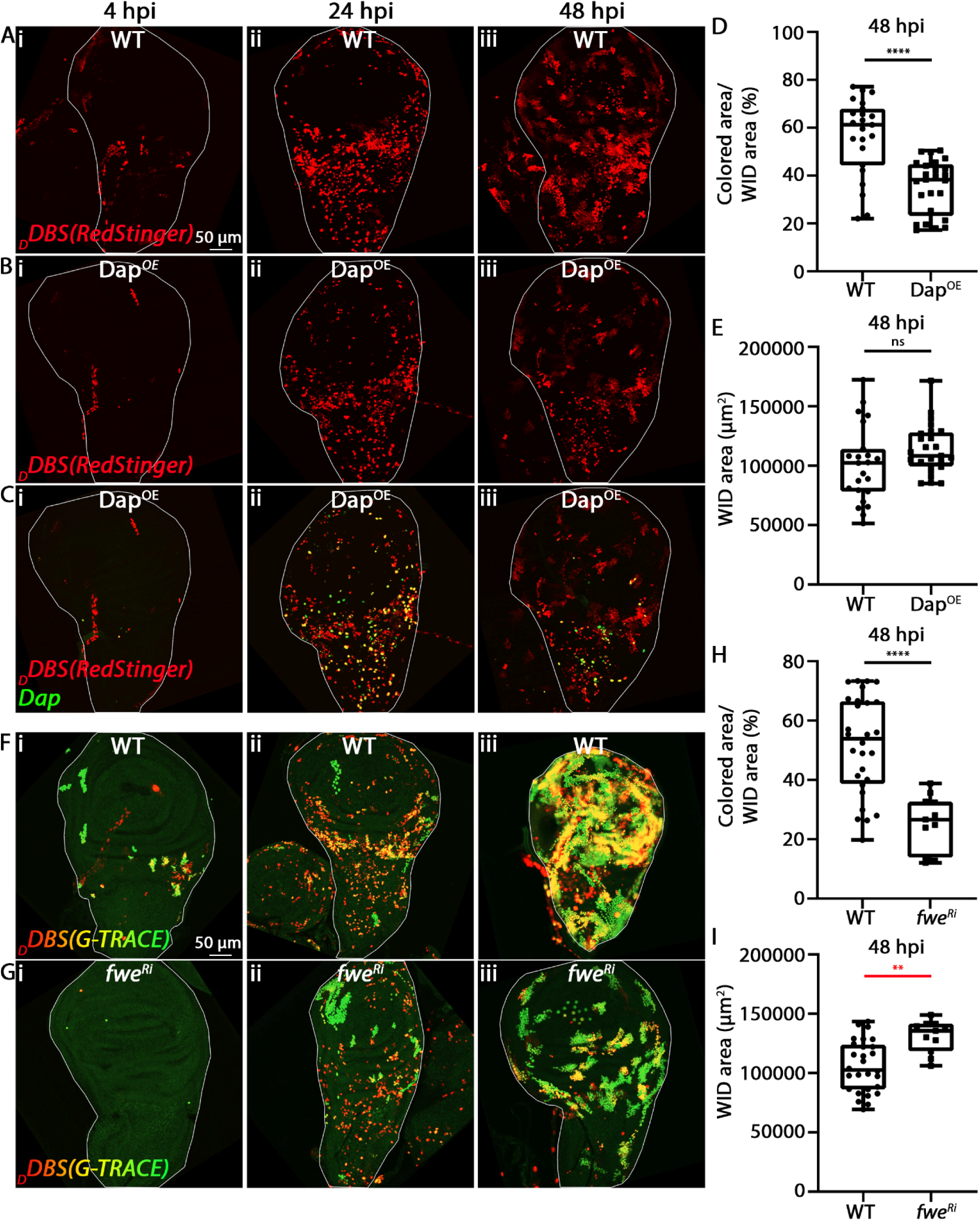
Attenuation of cell cycle or fitness in DARE cells leads to overproliferation of NARE cells. (A-C) Irradiated WIDs containing the *_D_DBS* reporter, *QUAS-Gal4* adapter, and *UAS-RedStinger*, with (B and C) or without (A) a *UAS*-*Dap-GFP* transgenic construct, were monitored at 4, 24, and 48 hpi. This transgenic combination enables simultaneous detection of DARE cells via RedStinger fluorescence (red) and overexpression of the cycle inhibitor Dacapo (Dap^OE^) in DARE cells. The WIDs in (B) and (C) are identical, with the distinction that (C) includes the green channel for detection of the DARE cells that overexpressed the Dacapo-GFP. Note the significant reduction at 48 hpi in both the number and size of DARE cell clones following the overexpression of Dacapo. Additionally, the few DARE cells that formed small clones exhibit no green fluorescence, indicating variability in the expression levels of the Dap transgene among different DARE cells. (D) Quantification of the number of DARE cells and their derivative daughter cells in 48 hpi WIDs, as represented in (A and B), is shown and presented as in Figure 3E. (E) The graph depicts the total size of the 48 hpi WIDs shown in (A and B), presented as the total area in square micrometers (µm²). The restoration of normal WID size at 48 hpi, despite the dramatic decrease in DARE cell proliferation, indicates overproliferation of NARE cells. (F and G) Irradiated WIDs expressing the transgenic combination described in Figure 3A, either without (F) or with a *UAS-fwe^Ri^*transgene (G), were monitored at 4, 24, and 48 hpi. While no detectable effects were observed at 4 and 24 hpi following knockdown of the *flower* (*fwe*) gene in DARE cells, a significant reduction in both the size and number of DARE cell clones was evident at 48 hpi. (H) Quantification of the number of DARE cells and their derivative daughter cells in 48 hpi WIDs, as represented in (F and G), is shown and presented as in Figure 3E. (I) The graph depicts the total size of the 48 hpi WIDs shown in (F and G), presented as the total area in square micrometers (µm²). Note that “labeling” DARE cells as less fit by knocking down *flower* in these cells results in increased proliferation of NARE cells and uncontrolled overgrowth of the WID. For the graphs in (D, E, H and I), *p* values were calculated using an unpaired Student’s t-test, two-tailed distribution. The graphs were generated and presented as in Figure 1B. \*\**p* < 0.01; \*\*\*\**p* < 0.0001; ns (non-significant).

The idea that DARE cell proliferation rate impacts NARE cell proliferation suggests that maintaining a proper balance between these cell populations may involve cell competition mechanisms, which typically monitor and regulate cell fitness during development and tissue homeostasis by eliminating less competitive cells. ^108–114^ To explore this possibility, we disrupted the process that differentiates between fit ‘winner’ cells and unfit ‘loser’ cells by knocking down the *flower* (*fwe*) gene in the DARE cells. Fwe is a conserved cell membrane protein essential for designating cells as winners or losers, and downregulation of *fwe* in WID cell clones surrounded by WT cells has been reported to trigger apoptosis in these clones. ^115,116^ While knockdown of *fwe* in DARE cells of irradiated WIDs with the *_D_DBS*/G-TRACE systems had no effect on the numbers of the DARE cells and derived clones at 4 and 24 hpi, a significant reduction in DARE cell-derived clones was observed at 48 hpi (Figures 7F-7H). Furthermore, at 48 hpi, the size of the *fwe* knockdown WIDs not only remained unchanged but was also significantly larger compared to their WT counterparts (Figures 7F, 7G, and 7I). This indicates that *fwe*-deficient DARE cells were outcompeted by neighboring NARE cells, and suggests that disrupting the proliferative homeostasis between these two cell populations can lead to unregulated tissue regrowth. Taken together, these findings highlight that proper compensatory growth of irradiated WIDs depends on a precise balance between DARE and NARE cell proliferation, which is regulated through the modulation of DARE cell proliferation.

## Discussion

In this study, we investigated the mechanisms underlying epithelial tissue regeneration by compensatory proliferation following high-dose IR. By employing a delayed genetic reporter for the activity of the initiator caspase Dronc, we identified a population of apoptosis-resistant epithelial cells - referred to as DARE cells - that are dispersed throughout the *Drosophila* imaginal discs. These cells largely emerge at 24 hpi and undergo extensive proliferation during the subsequent 24 hours.

Regenerated imaginal discs at 48 hpi also contain another population of cells, termed NARE cells, which do not activate the reporter. Although this suggests that NARE cells, unlike DARE cells, do not activate Dronc, it is also possible that they lack some of the transgenic elements needed for DARE cell detection, potentially due to mitotic recombination triggered by IR-induced double-strand DNA breaks. However, our findings, which show that DARE and NARE cells exhibit distinct proliferative characteristics, support the idea that NARE cells are largely a separate population of apoptosis-resistant epithelial cells that, along with the DARE cells, contribute to the regeneration of the imaginal disc.

In apoptosis-sensitive cells of the imaginal disc, Dronc activation on the Dark apoptosome leads to lethal activation of effector caspases and subsequent apoptosis. ^29^ In contrast, according to our model illustrated in Figure 8, Dark-independent activation of Dronc in DARE cells does not reach a lethal threshold of effector caspase activation due to its interaction with Myo1D. Consequently, DARE cells proliferate in a manner that depends on cell autonomous function of both Dronc and TNFR >JNK signaling, forming cell clones that replenish the imaginal disc. However, while Dronc signaling in DARE cells triggers NARE cell proliferation in a non-cell-autonomous manner, TNFR>JNK signaling does not. Proper tissue regeneration relies on maintaining proliferative homeostasis between DARE and NARE cell populations; disruption of this balance - e.g., by altering the TNFR>JNK signaling in the DARE cells - can result in one population outcompeting the other, which can lead to abnormal compensatory growth of the tissue.

**Figure 8.**
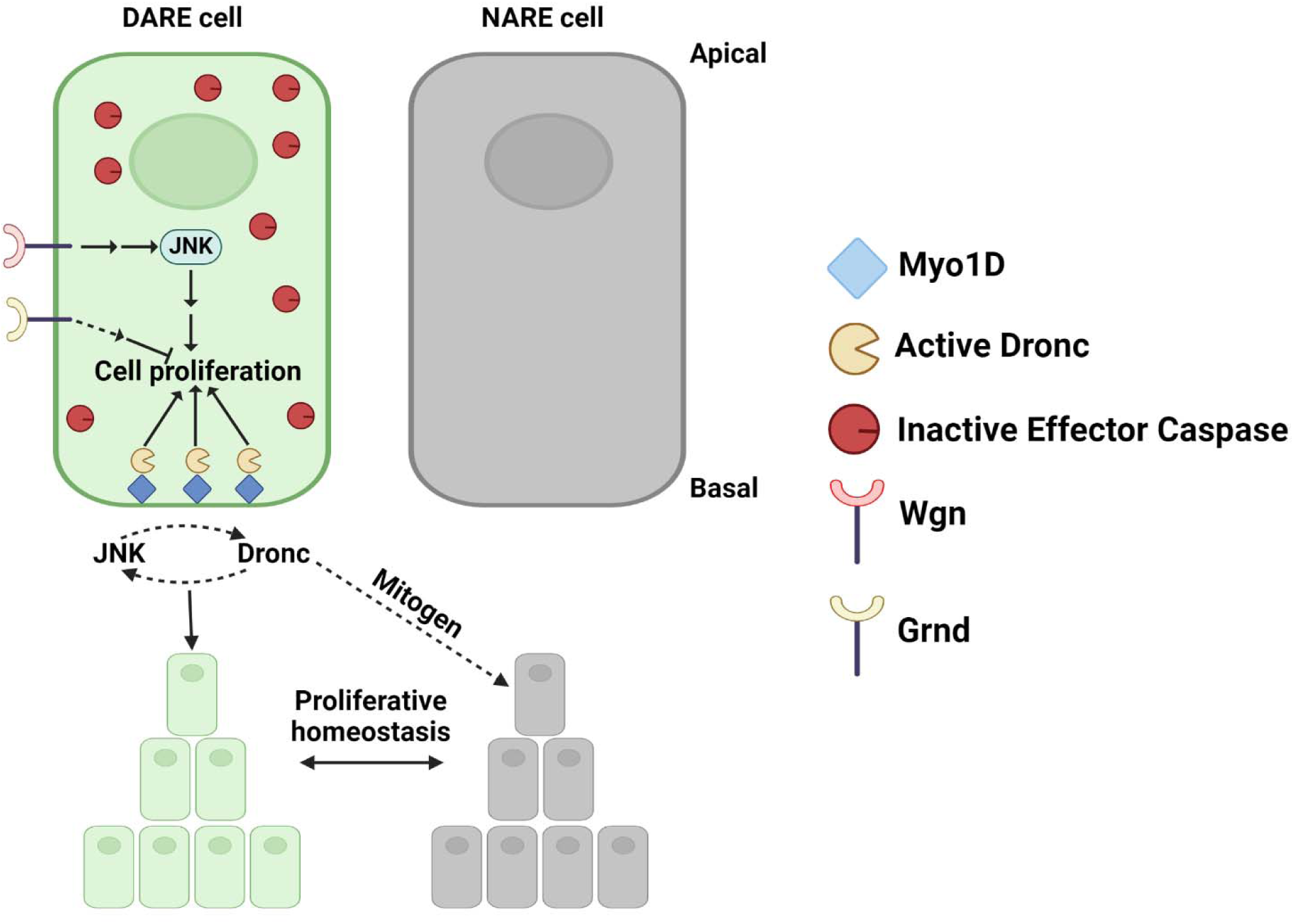
An integrated model of WID regeneration through compensatory proliferation following irradiation. The illustration at the top depicts two adjacent apoptosis-resistant epithelial cells: one from the DARE cell population (left) and one from the NARE cell population (right). At the bottom, the cell clones derived from the two cells illustrated at the top are shown. The signaling pathways in DARE cells are illustrated at the top left, with arrows representing activation and a T-bar indicating inhibition. The interplay between DARE and NARE cells is depicted at the bottom. A detailed explanation of the model can be found in the main text.

The mechanism by which Dronc activation occurs independently of the Dark apoptosome remains unclear. Furthermore, it is still uncertain whether such a mechanism is also linked to the ability of cells to activate Dronc while remaining alive. Dark-independent Dronc activation has been reported in other systems. For instance, Tango7 modulates Dronc levels and activity independent of Dark by sequestering it to the cell cortex during some non-apoptotic processes. ^91,117,118^ Additionally, Dronc has been shown to interact with Myo1D in ‘undead’ salivary gland and imaginal disc cells during AiP, as well as in enterocytes of the adult midgut. ^68,69^ This interaction sequesters Dronc to the basal side of the plasma membrane, where it exhibits minimal co-localization with Dark, which mainly resides in the cytoplasm, hence suggesting that Dronc is largely separated from Dark in these cellular contexts.^69^ Importantly, we demonstrate that this sequestration is crucial for preventing unwanted effector caspase activation and apoptosis of the DARE cells, providing proof of a mechanism by which normal cells can activate Dronc while remaining alive. Interestingly, MYO1D has been implicated in tumor growth and an increased risk of malignancy by anchoring EGFR family members to the plasma membrane, suggesting that this motor protein plays a conserved role in the membranal sequestration of proteins associated with cell survival and growth. ^119,120^

Although AiP is a non-physiological process, this study reveals significant overlaps between the mechanisms of AiP and compensatory proliferation. A key similarity is the central role of Dronc in driving cell proliferation in both contexts. ^38,64,65,68,69,71,75,76,95,121^ Additionally, as seen in DARE cells, both AiP and compensatory proliferation involve the JNK signaling pathway and TNFR. ^63,68,70,73,93,95,121,122^ Notably, Myo1D’s role in AiP and compensatory proliferation is particularly intriguing. While Myo1D has been reported to sequester Dronc to the plasma membrane during AiP - potentially enhancing Dronc’s non-cell autonomous proliferation effects - ^68–70^ our study suggests an additional critical function. The Myo1D-Dronc interaction also prevents excessive effector caspase activation and cell death in DARE cells. Thus, Myo1D’s influence extends beyond promoting cell proliferation to also safeguarding against cell death. Interestingly, the detection of Dronc at the plasma membrane in ‘undead’ cells - cells that would typically be destined for death - suggests that even in the context of cell death, Dronc may be positioned in a way that facilitates non-cell-autonomous proliferation through the AiP mechanism. This implies that dying cells could play a role in tissue regeneration by contributing to the overall regenerative process.

Regeneration and overgrowth of WIDs following IR or tissue injury have been previously attributed to epithelial cells that activate effector caspases but escape apoptosis, unlike their neighboring cells.^62,123,124^ It has been proposed that WIDs contain specific areas with apoptosis-resistant stem-like cells that act as reservoirs for tissue regeneration. ^62,123^ However, our findings indicate that, at 24 hpi, DARE cells are uniformly dispersed throughout the WID, with minimal clustering of DARE cell clones just below the pouch area. We demonstrate that the initiator caspase Dronc, rather than effector caspases, play a critical role in tissue regeneration. A plausible explanation for these discrepancies is that the CasExpress reporter used in previous studies might not be specific to effector caspase activity and could also be cleaved by Dronc. Consequently, these studies may have detected some Dronc-positive, effector caspase-negative cells (i.e., DARE cells) using the CasExpress reporter. Supporting this, it has been shown that overexpression of P35, which potently inhibits effector caspases but not Dronc, did not completely block CasExpress reporter activation in the eye imaginal disc. ^125^ This suggests that Dronc might contribute to the reporter’s activation. The CasExpress reporter includes a caspase-3-binding and cleavage domain from Diap1 with a DQVD cleavage site. ^125^ Although Dronc cleaves at TQTE site in Dronc and TETD site in Drice and Dcp-1, scanning synthetic combinatorial libraries of tetrapeptides with aspartate at the P1 position showed that Dronc has a strong preference for valine at the P2 position. A broader range of amino acids is tolerated at the P3 and P4 positions, including glutamine and aspartate, respectively. ^36^ Therefore, Dronc has the potential to cleave at the DQVD site, which could lead to the activation of the CasExpress reporter.

Understanding the mechanisms underlying compensatory proliferation following ionizing irradiation is crucial for both advancing our knowledge of tissue regeneration and addressing its connection to cancer. ^57^ Key characteristics of cancer cells are their ability to evade apoptosis and proliferate uncontrollably. ^126^ Indeed, several key factors associated with compensatory proliferation identified in the current study have previously been shown to promote tumor formation when dysregulated. Myo1D was found to be essential for neoplastic tumor growth and invasion in *scrib*^-/-^*Ras^V^*^12^ cells, likely by promoting the localization of Dronc to the plasma membrane. ^69^ Furthermore, expression of both MYO1D and MYO7A were linked to tumor growth and increased risk of malignancy. ^119,120,127^ The role of JNK in promoting tumor formation has also been demonstrated across various cancer models, ^128–132^ including in *Drosophila*. ^63,92,96,133–137^ Given that most current anticancer agents, including IR, induce apoptotic cell death ^138–142^ and that approximately two-thirds of cancer patients undergo radiotherapy, ^143–146^ elucidating how certain cells resist radiation-induced apoptosis and subsequently proliferate could enhance our understanding of cancer recurrence after therapy ^147–150^ and lead to the development of more precise and effective cancer treatments.

## Methods

### Fly strains

For a comprehensive list of the fly lines used in this study, please refer to the key resources table. The control lines used in the experiments share the same genetic background as the tested lines, except for the specific conditions being tested. These control lines are typically referred to as wild type (WT). The flies were reared at 25 °C.

### Generation of the constitutively active Drice (*actDrice*) construct

A 935 bp fragment encoding the large and small subunits of Drice, excluding the prodomain, was PCR amplified from a full-length *drice* cDNA clone using the following primers: a forward primer, 5’-AATGCGGCCGCATGGCCCTGGGCTCCGTGGGAT-3’, which includes an ATG transcription start site and a NotI restriction site, and a reverse primer, 5’-CCAAGGTACCGCCAGCGGCCC-TCAAACCC-3’, which includes a KpnI restriction site. This fragment was then directionally subcloned into the NotI and KpnI sites of the *pUASTattB* plasmid, resulting in the *pUAST-NoPro-drice* plasmid. A 66 bp DNA fragment of the viral P2A sequence was PCR amplified from a plasmid provided by Prof. Yosef Shaul (in our department), using the following primers: a forward primer, 5-’ ACCATGCAGCGTTCTCAGACGGAAACCGATGGAAGCGGAGCTACTAACTT-3’, and a reverse primer, 5’-TGGAATCTTGTAGCTCATCGAGGAGTCGCCAGGTCCAGGGTTCTCCTC-3’. Both primers included *drice* overhang sequences, which facilitated the insertion of the amplified P2A fragment into the *pUAST-NoPro-drice* plasmid, between the large and small subunits of Drice, using restriction-free cloning. The P2A fragment was inserted at position 609 of the *NoPro-drice* sequence, immediately following the TETD tetrapeptide at the end of the large subunit.

Transgenic flies were generated via φC31-mediated site-specific transgenesis, with injections performed by BestGene Inc. (Chino Hills, CA, USA). The transgene was inserted into the attP40 site on the second chromosome.

### Immunostaining procedures and imaging

All *Drosophila* larvae were irradiated with a dose of 20 Gy X-rays using the XRAD 320 X-ray unit from Precision X-ray (USA), except for the experiment shown in Figure 3G, where a dose of 10 Gy X-rays was used to facilitate recovery of adult flies.

Imaginal discs were dissected from third instar larvae either before irradiation or at various time points afterward. The order of imaginal disc dissections by genotype was randomized for each experiment. After dissection, the imaginal discs were fixed in 4% paraformaldehyde (PFA) in PBS for 20 minutes at room temperature, then washed three times in PBX (PBS with 0.1% Triton X-100) for 10 minutes each. If no additional staining was needed, the discs were mounted and examined using a Dragonfly 505 spinning disk confocal system (Andor Technology PLC). Images were captured with a Sona 6 sCMOS camera mounted on a Leica DMi8 inverted microscope.

Immunostaining of the imaginal discs was performed on the washed fixed samples. The discs were first incubated in a blocking solution of 1% BSA in PBS for 1 hour at room temperature, then incubated overnight with a primary antibody diluted in 1% PBS/BSA at 4 °C. The samples were then washed in PBX, incubated with the secondary antibody for 1 hour at room temperature, washed again in PBX, and finally mounted. In experiments involving multiple time points, samples were stained and mounted together at the latest time point for comparison.

To visualize the adult wing, newly eclosed adults with unfurled wings were collected and kept at room temperature until the wings fully expanded. The wings were then collected within a few minutes of unfurling, mounted, and imaged immediately.

### Antibodies

Primary antibodies used in this study included rabbit polyclonal anti-cleaved human PARP (diluted 1:500, Ab2317; Abcam), rabbit polyclonal anti-pHH3 (diluted 1:300, 06-570; Millipore), and rabbit anti-cleaved Dcp-1 (diluted 1:200, 9578; Cell Signaling). Secondary antibodies were sourced from Jackson ImmunoResearch and were used at a dilution of 1:200.

### TUNEL labeling

WIDs were fixed in 4% PFA for 20 minutes, washed twice in PBS (2 × 5 minutes), washed twice (2 × 10 minutes) in 1xBSS (5 x BSS: 270 mM NaCl, 200 mM KCl, 37 mM MgSO4, 12 mM CaCl2.2H2O, 24 mM tricine, 1.8% glucose, and 8.5% sucrose), followed by two washes (2 × 5 minutes) in PBTw (0.1% Tween 20 in PBS). The samples were then refixed in 4% PFA for 20 minutes, washed five times (5 × 5 minutes) in PBTw, incubated in equilibration buffer (ApopTag kit; Millipore) for 1 hour, followed by overnight incubation with the TdT enzyme in reaction buffer (ratio 3:7; ApopTag kit) at 37 °C. The reaction was stopped by replacing the reaction mix with stop buffer (diluted 1:34 in dH2O; ApopTag kit) and incubation for 3-4 hours at 37 °C. The samples were then washed three times (3 × 5 minutes) in PBTw, blocked in BTN solution (1xBSS, 0.3% Triton X-100, and 5% normal goat serum) for 1 hour at room-temperature, and incubated overnight in the dark with anti-digoxigenin antibody solution (diluted 47:53 in blocking solution; ApopTag kit) at 4 °C. Samples were then washed four times (4 × 20 minutes) in 1xBSS, and mounted in Fluoromount-G (SouthernBiotech).

### Preparation and imaging of *Drosophila* eyes

Male fly heads were detached from the bodies using a needle and then further cut to separate the two eyes. Images were captured using a stereo microscope (SZX16; Olympus) equipped with a DP28 camera (Olympus).

### Quantification of imaging data

WIDs were imaged by capturing optical Z-slices at 40-micron intervals, resulting in approximately 90 Z-stacks per WID. Post-acquisition image processing and analyses were conducted using Fiji. ^151^

In all images except for the blue graph in Figure S2, quantifying the different fluorescent signals expressed in the WIDs under various genetic conditions, was done by a semi-automated ImageJ Fiji script developed to enable precise measurement of regions of interest. In brief, a 2D maximal intensity projection of the Z-stack for each imaged WID was generated. Each fluorescent channel was then thresholded and masked under careful supervision to prevent misclassification. Thresholding for the entire imaged WID was performed using a combination of all fluorescent channels. The masked images were then used to define overlaying and divergent regions of interest for all relevant imaged channels. Measurements of the mean intensity and area of the resulting regions of interest were conducted on the corresponding original 2D maximal intensity projection images for each fluorescent channel. All regions of interest and processed images were documented and saved. The results were exported as .csv files for further statistical analysis.

For the quantifications in Figure S2 (blue graph), images were analyzed using the open-source software Fiji. ^151^ For each image, a user-defined outline of the WID of interest was applied, along with an intensity threshold for each dye. This allowed for the quantification of the areas where each dye was expressed and the measurement of the overlap between these areas in each slice.

Both scripts will be available in GitHub.

### Data presentation, statistical analysis, and reproducibility

Illustrations in Figures 1D, 1E, 2A, 3A, 4G, 5A, 6A, 8, and S1A were created using BioRender.com. All graphs and statistical analyses in this manuscript were generated using the GraphPad Prism software version 9.5.1 for Windows (GraphPad Software, San Diego, California USA, www.graphpad.com). The specific statistical tests used to determine significance between experimental groups, as well as details on experimental reproducibility, are provided in the relevant figure legends. Significance is indicated by asterisks as follows: \**p* < 0.05, \*\**p* < 0.01, \*\*\**p* < 0.001 and \*\*\*\**p* < 0.0001. All experiments were conducted with at least two biological replicates. Sample size was determined based on standard practices within the laboratory and the general field of study.

## key resources table

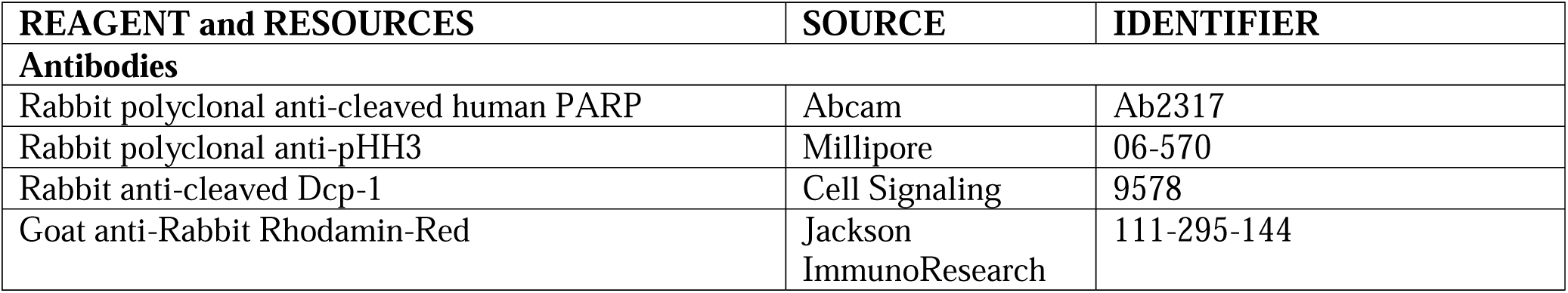
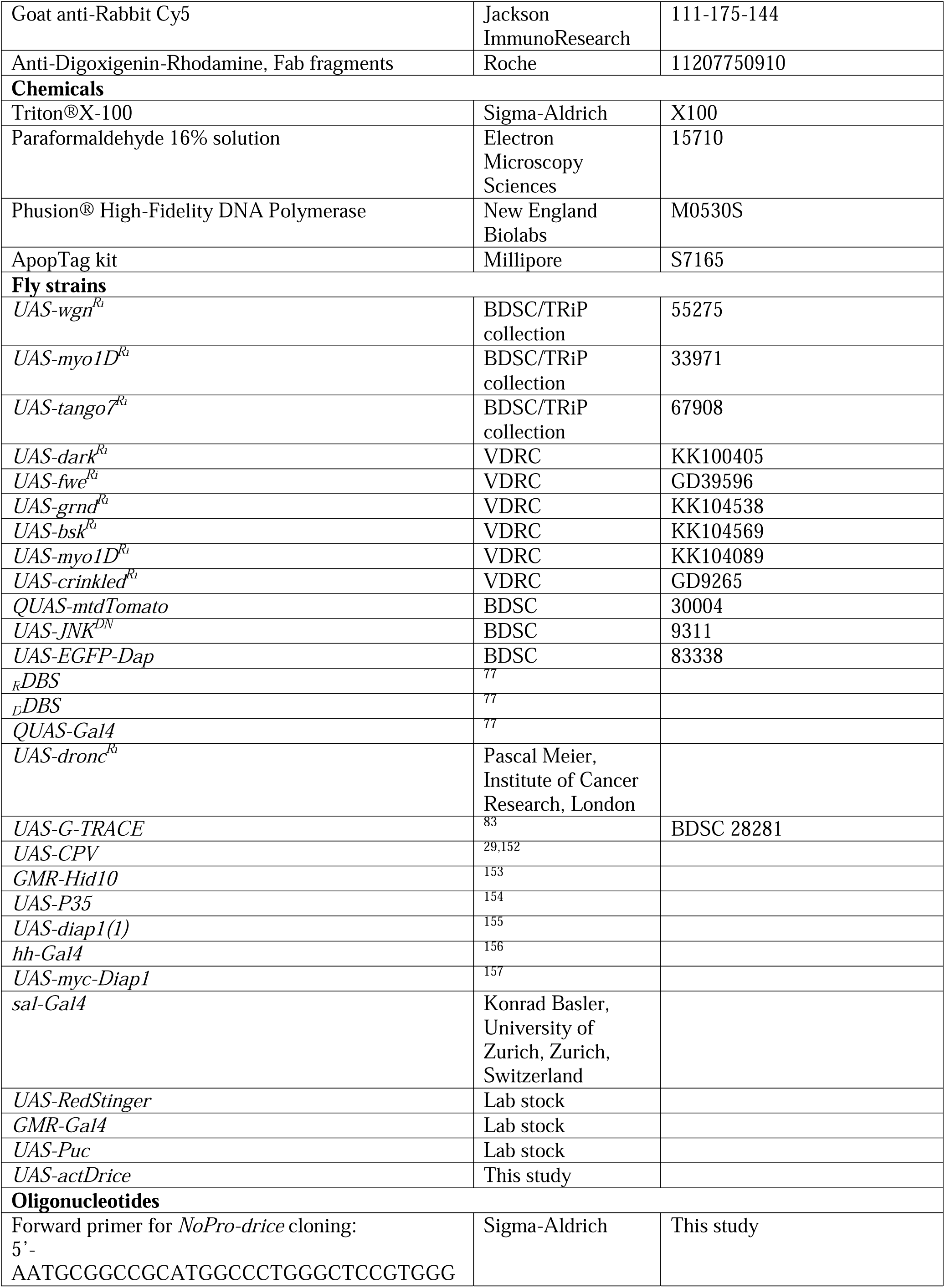
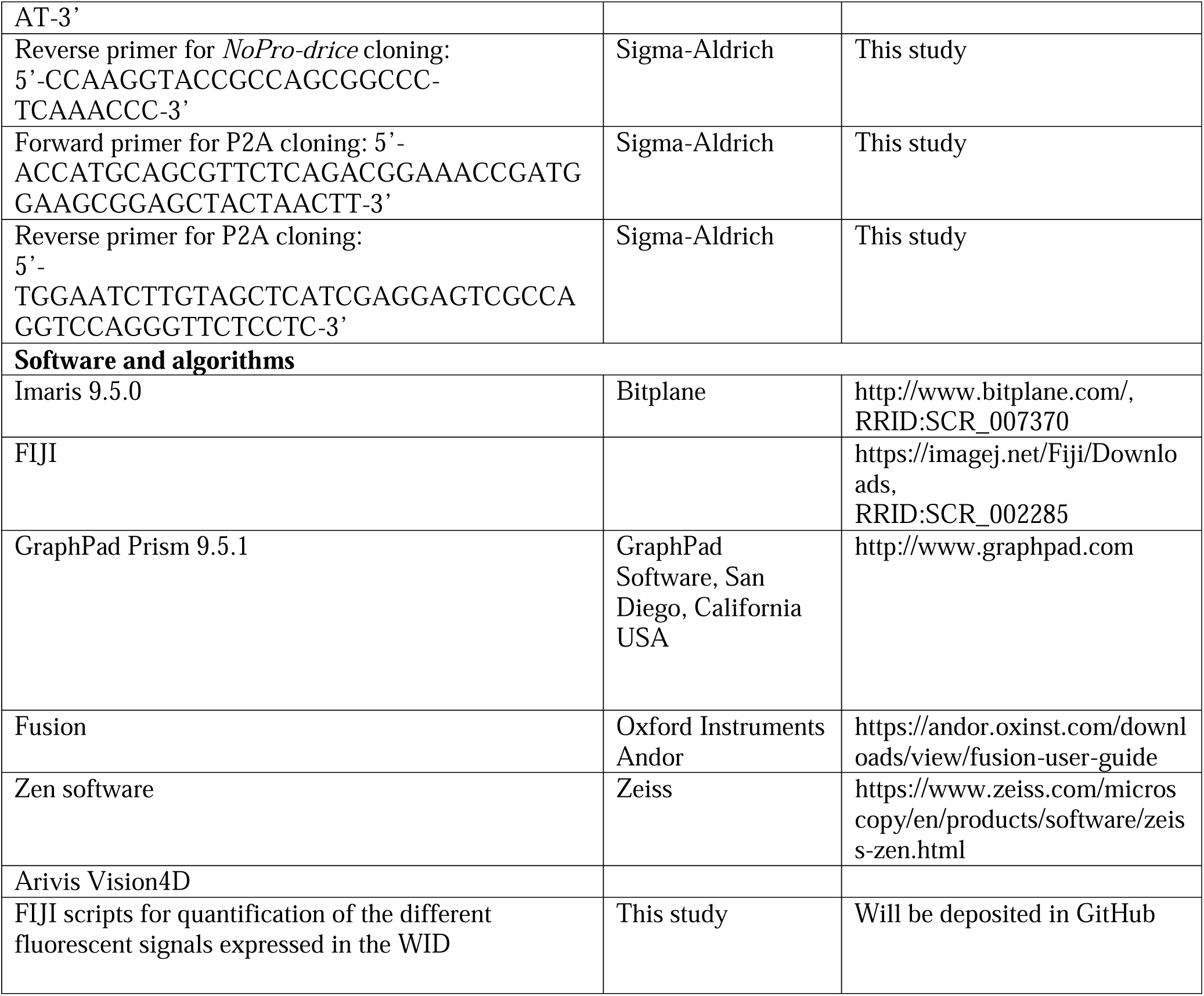

### Declaration of generative AI and AI-assisted technologies in the writing process

During the preparation of this work the author(s) used ChatGPT in order to improve readability and language of several sentences in this work. After using this tool/service, the author(s) reviewed and edited the content as needed and take(s) full responsibility for the content of the publication.

## Acknowledgments

We thank Pascal Meier, Oren Schuldiner, Talila Volk, the Vienna *Drosophila* Resource Center (VDRC), the Transgenic RNAi Project (TRiP), and the Bloomington *Drosophila* Stock Center for providing essential stocks and reagents. We also appreciate Ron Rotkopf for his assistance with statistical analysis, Anna Gorelick-Ashkenazi for providing the control image for the *CPV* reporter, and the Arama laboratory members for their encouragement and advice. Additionally, we acknowledge the BioRender website for its role in creating the illustrations used in this paper.

---

**Figure S1.**
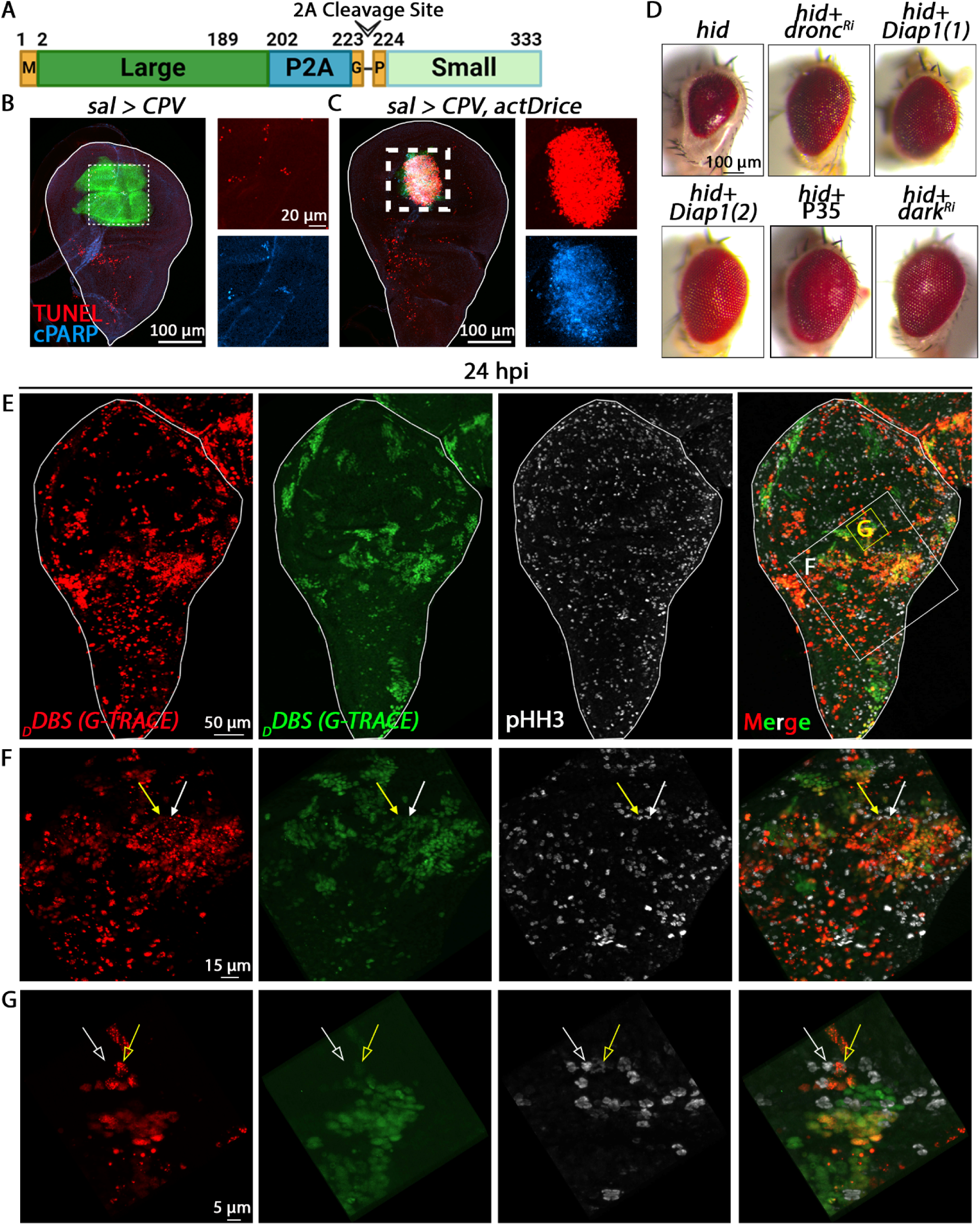
Regeneration of the WID after irradiation is mediated by proliferating DARE and NARE cells. (A) A schematic structure of the constitutively active Drice (actDrice) construct is illustrated, showing the first methionine inserted in place of the N-terminal prodomain, with the large and small subunits of Drice separated by the P2A ribosomal skipping peptide. (B and C) Representative WIDs from 3^rd^ instar larvae expressing the *CPV* reporter for effector caspase activity (see Figure 5A for *CPV* structure and mode of action) in the pouch region using the *sal-Gal4* driver are shown. The WIDs were immunostained with an anti-cleaved human PARP antibody (cPARP; blue) to reveal effector caspase activity and labeled with TUNEL to detect dying cells (red). The *sal-Gal4* expression domain is indicated in green due to the Venus fluorescence of the *CPV* reporter. Enlargements of the areas outlined by dashed squares are shown to the right of the WIDs, displaying the red and blue channels separately. Note the minimal effector caspase activation (cPARP) and apoptosis (TUNEL) in the control WID (B), compared to the excessive effector caspase activity and massive cell death observed with the expression of the actDrice construct (C). (D) Assessment of the effectiveness of the apoptosis-related transgenes used in this study for inhibiting apoptosis induced by ectopic expression of the pro-apoptotic protein Hid in *Drosophila* photoreceptor cells. Transgenesis experiments using the *Drosophila* adult compound eye are shown. The transgenes indicated above each panel were ectopically expressed in developing photoreceptors using the *GMR-Gal4* driver. Note that two independently generated Diap1 transgenes were used in this study: Diap1(1) in Figure 4E and Diap1(2) in Figure 4H. Representative eyes from newly eclosed males are displayed. Note the typical small eye phenotype resulting from *hid* expression, and the suppression of this phenotype when these transgenes are co-expressed. (E) Irradiated WID expressing the G-TRACE system in DARE cells, as described in Figure 3A, were immunostained with an anti-Phosphohistone H3 (pHH3) antibody to identify proliferating cells in the G to M phase. Shown is a representative WID at 24 hpi, with the red and green channels presented separately to distinguish between newly appearing DARE cells (red) and DARE cell clones (green). Note that the pHH3-positive cells are distributed fairly uniformly among both DARE (colored) and NARE (uncolored) cells. (F and G) Enlargements of the areas marked by squares in the WID shown in (E). Arrows in (F) denote two dividing DARE cells (marked by both red and green, as well as pHH3) located at the periphery of DARE cell clones. In (G), the arrows highlight two dividing cells: the yellow arrow points to a DARE cell (marked by red, green, and pHH3), while the white arrow points to a closely adjacent dividing NARE cell (marked by pHH3, but showing no fluorescence from the G-TRACE system).

**Figure S2.**
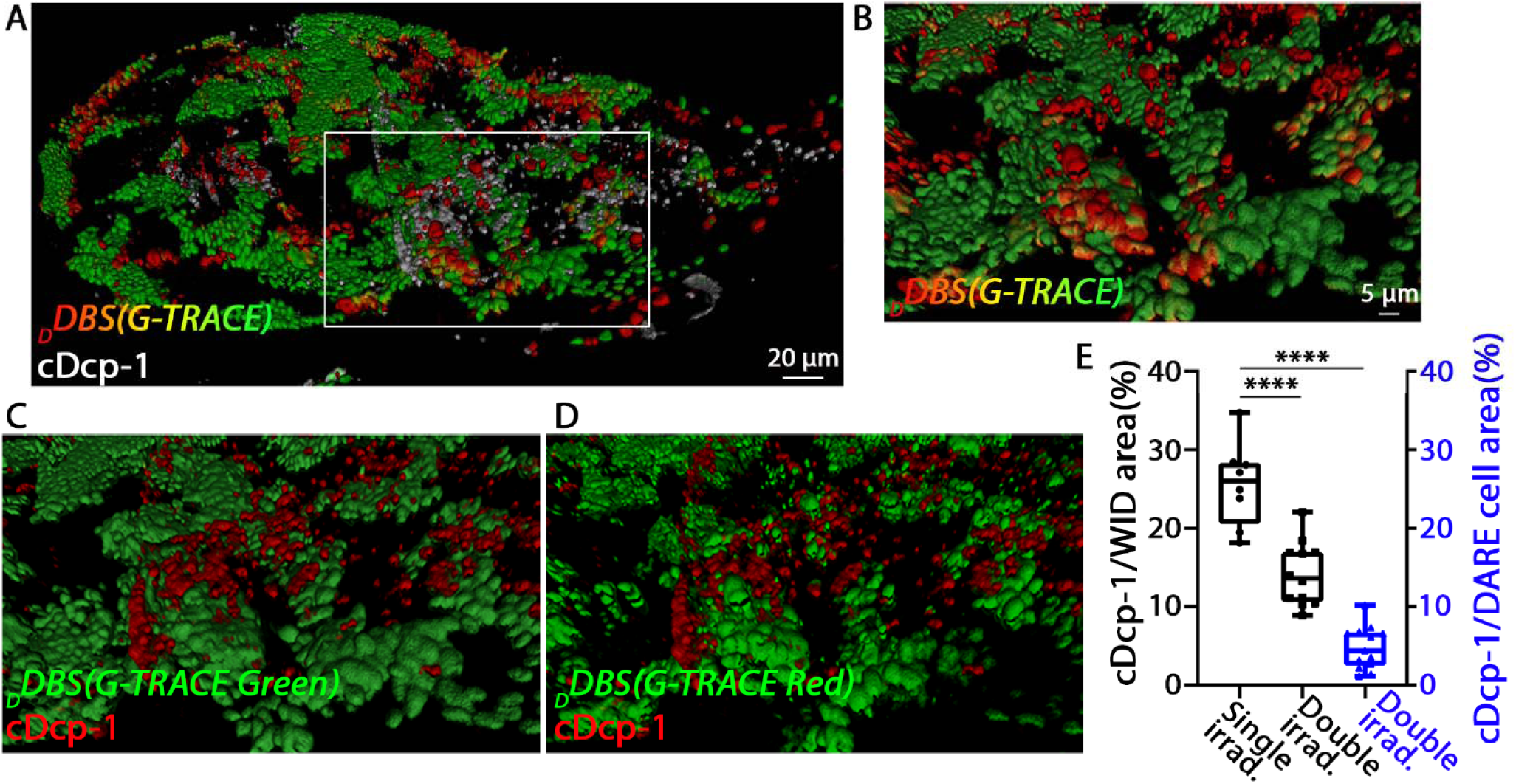
DARE cell-derived daughter clones are resistant to re-irradiation-induced apoptosis. (A) A representative irradiated WID expressing the *_D_DBS*/G-TRACE systems (as described in Figure 3A) was re-irradiated at 48 hours post-initial irradiation, then dissected 4 hours later and immunostained with an anti-cDcp-1 antibody to identify dying cells (cDcp-1; white/gray). The DARE cells and derived clones detected with the G-TRACE system are indicated in red and green. Overlapping red and green cells are yellow. (B-D) Enlargements of the area outlined by the white rectangle in (A) show the overlap between the three channels more clearly. To facilitate this, each image displays only two channels at a time, represented in green and red. Specifically: the classical colors of the G-TRACE system is shown in (B), DARE cell clones in green and cDcp-1 expression in red are depicted in (C), and early DARE cells in green with cDcp-1 expression in red are illustrated in (D). Notably, in (B), the overlap between channels is visible as yellow in early DARE cells, whereas in (C) and (D), there is minimal overlap between cDcp-1 and DARE cells. (E) The black graphs depict the quantification of dying cells in WIDs as shown in Figure 1B, measured 4 hours after the first and second irradiations. The data is presented as the percentage of the WID area occupied by dying cells, calculated by dividing the area of dying cells by the total WID area and multiplying by 100. Note that the average area occupied by dying cells decreases from 25.6% four hours after the first irradiation (Single irrad.) to 13.4% four hours after the second irradiation (Double irrad.), representing roughly a 50% reduction. The blue graph illustrates the quantification of DARE cells and their derived daughter cells expressing cDcp-1, expressed as a percentage. This percentage is calculated by dividing the area occupied by DARE cells and their clones that express cDcp-1 by the total area of DARE cells and clones, and then multiplying by 100 to represent the proportion undergoing apoptosis. To ensure accuracy, 20 optical Z-slices were analyzed per WID to measure the overlaps. Note that the average likelihood of a DARE cell undergoing apoptosis 4 hours after the second irradiation is 4.8%. This is approximately five times lower than the 25.6% observed 4 hours after the first irradiation (see Figure 1B). *p* values were calculated using an unpaired Student’s t-test, two-tailed distribution. The graphs were generated and presented as in Figure 1B. \*\*\*\**p* < 0.0001.

**Figure S3.**
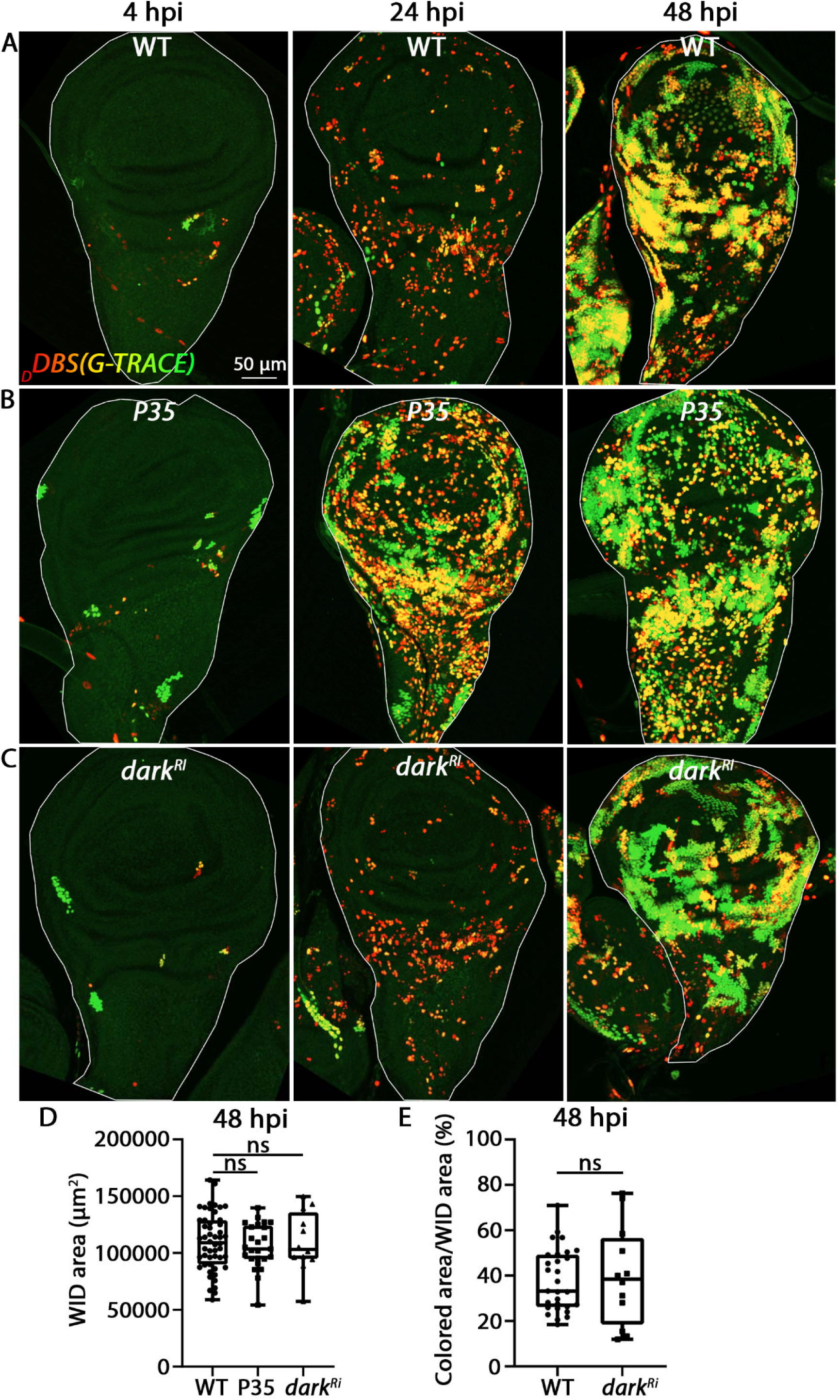
Effector caspases and the Apoptosome adapter Dark are not required for tissue regeneration by compensatory proliferation. (A-C) Irradiated WIDs expressing the transgenic combination described in Figure 3A were monitored at 4, 24, and 48 hpi. The conditions include: no additional transgene (A), a *UAS-P35* transgene (B), or a *UAS-dark^Ri^* transgene (C). Both conditions had no effect on DARE cell proliferation or on the number and size of DARE cell clones at 48 hpi (green). However, ectopic expression of P35 blocks the natural apoptosis of many DARE cells due to its inhibitory effect on effector caspase activity, leading to their accumulation at 24 and 48 hpi (single red and yellow cells). |(D) The graphs depict the total size of the 48 hpi WIDs shown in (A-C), presented as the total area in square micrometers (µm²). Note the restoration to normal size at 48 hpi compared to the control WT WID counterparts. (E) Quantification of the number of DARE cells and their derivative daughter cells in 48 hpi WIDs, as represented in (A and C), is shown and presented as in Figure 3E. For the graphs in (D and E), *p* values were calculated using an unpaired Student’s t-test, two-tailed distribution. The graphs were generated and presented as in Figure 1B. ns, non-significant.

**Figure S4.**
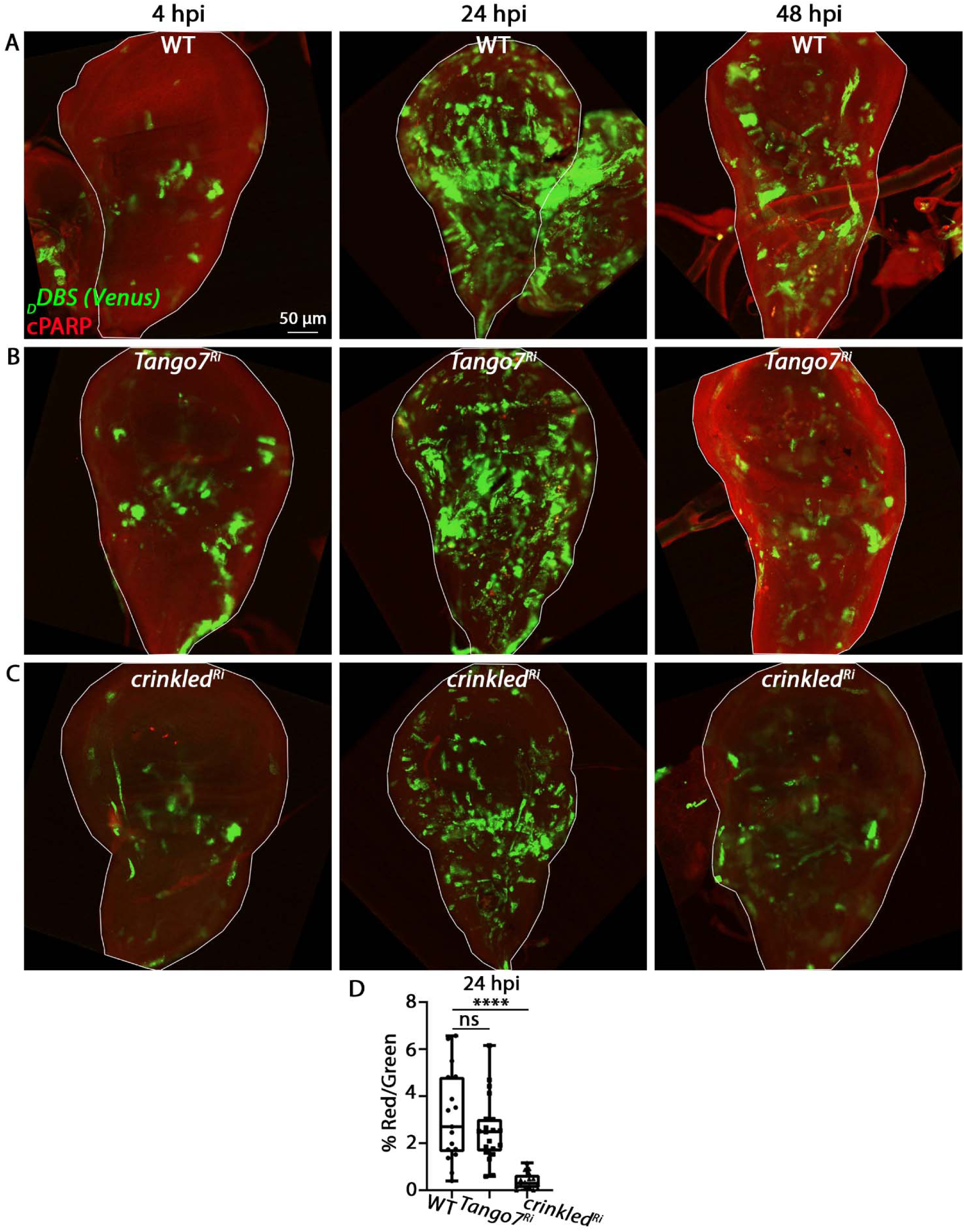
The Dronc-interacting protein Crinkled, but not Tango7, acts as a positive regulator of effector caspase activation in DARE cells. (A-C) Irradiated WIDs expressing the transgenic combination described in Figure 5A, either without (A) or with a *UAS-tango7^Ri^*transgene (B) or a *UAS-crinkled^Ri^* transgene (C), were immunostained for cPARP and monitored at 4, 24, and 48 hpi. DARE cells are visualized by the Venus fluorescence of the *CPV* (green) and/or by cPARP (red) after its cleavage by the effector caspases. (D) Quantification of the number of DARE cells that activated effector caspases (i.e., dying cells; cPARP) in WIDs represented in (A-C). The graph represents the area occupied by cPARP-expressing DARE cells (red) divided by the area covered by DARE cells without effector caspase activity (green), multiplied by 100 to express the percentage of DARE cells exhibiting effector caspase activity. Note the significant decrease of effector caspase activity in *crinkled^Ri^*-expressing DARE cells. The graph was generated and presented as in Figure 5E. \*\*\*\**p* < 0.0001; ns (non-significant).

**Figure S5.**
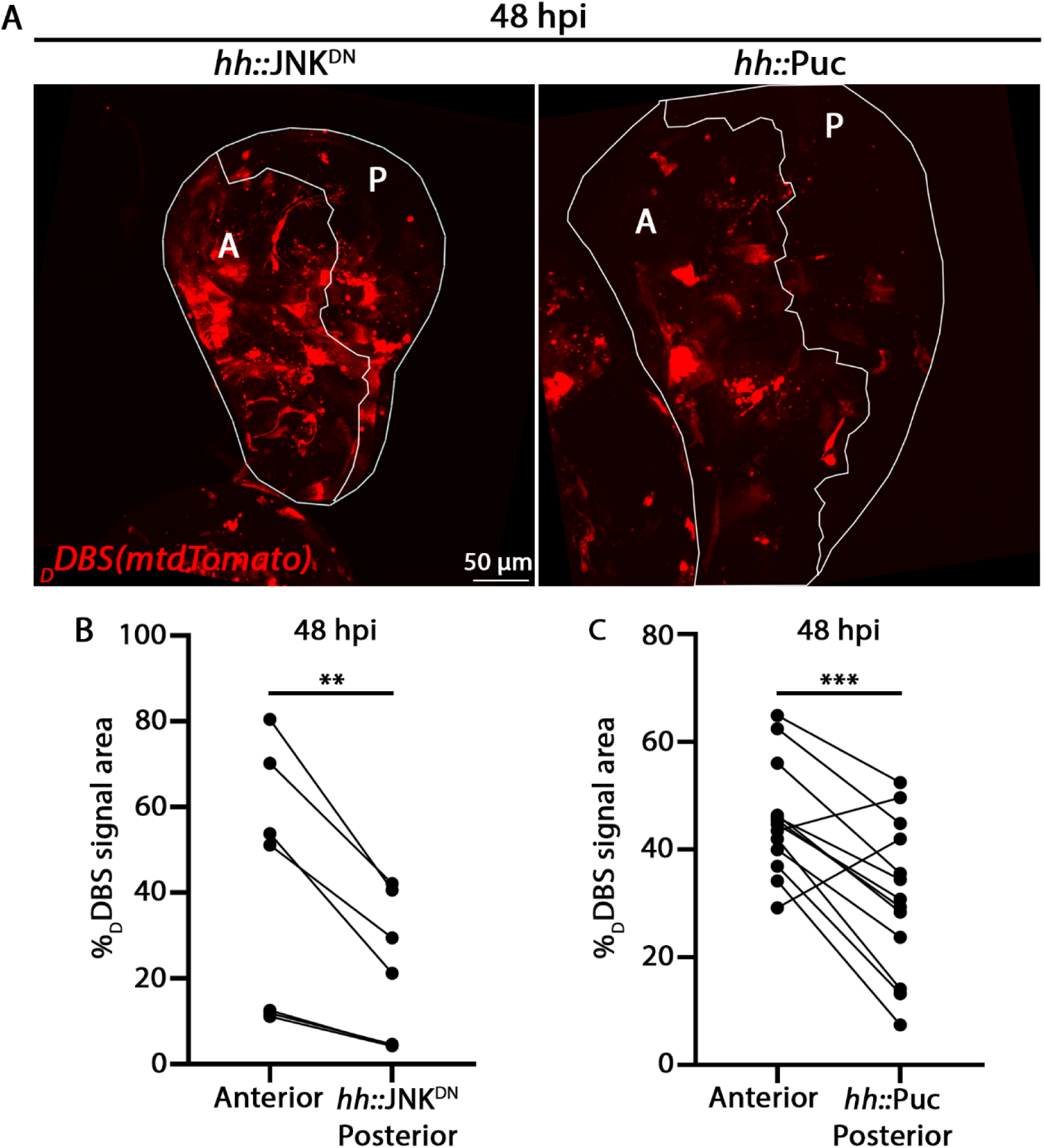
JNK in the DARE cells is required for DARE cell proliferation. (A) Irradiated WIDs of the genotype described in the scheme shown in Figure 4G, expressing exclusively in the posterior compartment either a dominant-negative JNK transgene (*UAS-jnk^DN^*) or the JNK inhibitor phosphatase Puckered (*UAS-Puc*). Note the dramatic decrease in DARE cell clones observed in the posterior compartments of 48 hpi WIDs with both conditions, while the unmanipulated anterior compartments show no such decrease (see also a control WID in Figure 4H). (B and C) Before-and-after presentation of individual values shown as the percentage of area occupied by DARE cell clones in the posterior and anterior compartments of the WIDs represented in (A). The graphs were generated and presented as in Figure 4I. For the graphs in (B and C), *p* values were calculated using a paired Student’s t-test, two-tailed distribution. \*\**p* < 0.01; \*\*\**p* < 0.001.

